# Fluoxetine promotes immunometabolic defenses to mediate host-pathogen cooperation during sepsis

**DOI:** 10.1101/2023.11.18.567681

**Authors:** Robert M. Gallant, Jessica M. Snyder, Janelle S. Ayres

## Abstract

Selective serotonin reuptake inhibitors (SSRIs) are some of the most prescribed drugs in the world. While they are used for their ability to increase serotonergic signaling in the brain, SSRIs are also known to have a broad range of effects beyond the brain, including immune and metabolic effects. Recent studies have demonstrated that SSRIs are protective in animal models and humans against several infections, including sepsis and COVID-19, however the mechanisms underlying this protection are largely unknown. Here we mechanistically link two previously described effects of the SSRI fluoxetine in mediating protection against sepsis. We show that fluoxetine-mediated protection is independent of peripheral serotonin, and instead increases levels of circulating IL-10. IL-10 is necessary for protection from sepsis-induced hypertriglyceridemia and cardiac triglyceride accumulation, allowing for metabolic reprogramming of the heart. Our work reveals a beneficial “off-target” effect of fluoxetine, and reveals a protective immunometabolic defense mechanism with therapeutic potential.

## Introduction

Host defense strategies against pathogens can be broadly categorized based on their effects on pathogen fitness (Schneider and Ayres, 2008). Antagonistic defenses protect the host by having a negative impact on pathogen fitness. They include both resistance defense mechanisms that kill pathogens, and avoidance mechanisms that prevent exposure to pathogens. In contrast, cooperative defenses facilitate host adaptation to the infected state, and protect the host while having a neutral to positive effect on pathogen fitness. They include both anti-virulence mechanisms that block host and microbial derived pathogenic signals, and disease tolerance mechanisms that limit host susceptibility to physiological dysfunction and damage (Ayres, 2020a).

Treatment strategies for infectious diseases will follow these same principles. Drug interventions can have antimicrobial actions via direct effects on the pathogen, such as antibiotics, or indirectly by heightening the host immune response to the infection. They can also promote host-pathogen cooperation by acting to neutralize/block pathogenic signals that can cause damage, for example as seen with the anti-inflammatory actions of steroids, or promote disease tolerance by limiting susceptibility to physiological dysfunction/damage, such as the use of fluid replacement or vasoconstrictors to raise blood pressure in critical illness. The majority of infectious disease treatments work to rid us of pathogens. However, for some infectious diseases, including sepsis, the host response to the infection does far more damage than the pathogen and thus anti-microbial based therapeutics are often insufficient for patient survival within these contexts (Cavaillon et al., 2020). New host-targeted therapeutic approaches that neutralize or detoxify host-derived pathogenic signals, and that protect from physiological dysfunction and damage in response to pathogenic signals, are needed.

Sepsis is defined as a dysregulated systemic inflammatory response that can lead to multi-organ failure and death (Hotchkiss et al., 2016). The most obvious way to target the host response in a septic patient is to suppress the inflammatory response. Indeed, numerous clinical trials have been performed with neutralizing antibodies that target pro-inflammatory cytokines, however such approaches have been met with little success (Fisher, 1994; Fisher et al., 1996; Gotts and Matthay, 2016; Hotchkiss et al., 2016; Panacek et al., 2004). One explanation for why this strategy has been largely unsuccessful is that such strategies can immunocompromise the patient, making their primary infections more difficult to control and/or rendering them more susceptible to secondary infections. A second possible explanation is that timing is critical, and such interventions must be administered prior to damage occurring (Ayres, 2020b). Additionally, sepsis is a multi-factorial syndrome with many derangements in host physiology beyond immune over-activation including dysfunction of the metabolic response of the patient. Host directed therapeutics that can control the degree and duration of an immune response to allow pathogen killing but prevent the escalation of the response to a cytokine storm, and that also promote metabolic adaptation to the infected state, may offer more therapeutic benefit than strategies that focus only on blocking the pro-inflammatory response.

Selective serotonin reuptake inhibitors (SSRIs) are some of the most widely prescribed drugs with over 13% of adults using these antidepressant medications between 2015 and 2018 (Brody and Gu, 2020). SSRIs were originally developed in the 1970s for their ability to prevent serotonin reuptake in the synaptic cleft of serotonergic neurons (Wong et al., 1974). It is now recognized that SSRIs also have a wide range of peripheral effects including regulation of immune and metabolic processes (Olguner Eker et al., 2017; Szałach et al., 2019). Furthermore, SSRIs have been shown to protect against sepsis in animal models (Rosen et al., 2019) and improve outcomes in patients infected with SARS-CoV-2 (Reis et al., 2022). The mechanisms underlying these protective effects are unclear. SSRIs have been reported to have anti-inflammatory effects, which suggests they may protect against overwhelming inflammatory responses and cytokine storm (Durairaj et al., 2015; Tynan et al., 2012). They have also been reported to regulate aspects of systemic metabolism that are dysregulated during sepsis and other inflammatory states including lipid metabolism (Chiu et al., 2021; Pan et al., 2018; Rozenblit-Susan et al., 2016). It remains unclear whether the anti-inflammatory effects of SSRIs are somehow related to the drug’s effect on lipid metabolism or if these are independent effects of SSRIs that may regulate host survival.

Here, we examined how the SSRI, fluoxetine, regulates survival and disease progression in a mouse model of sepsis. We found that fluoxetine promotes both resistance and cooperative defenses in a serotonin-independent manner. Instead, fluoxetine increases circulating levels of IL-10, which in turn protects from sepsis induced hypertriglyceridemia and cardiac triglyceride accumulation, allowing metabolic reprogramming in the heart during infection. Our study mechanistically links the anti-inflammatory and metabolic effects of SSRIs, and demonstrates that fluoxetine can be used as a prophylactic to protect from sepsis induced lethality by orchestrating protective immunometabolic mechanisms which may be leveraged and further explored by future studies.

## Results

### Fluoxetine protects from sepsis-induced disease and mortality

To study the effects of fluoxetine during infection, we conducted a pre-treatment dose titration as shown in **Figure 1A** based on previously published chronic dosing regimens (Dos Santos et al., 2006; Gomes et al., 2009). Following this pre-treatment, we challenged mice intraperitoneally with our polymicrobial sepsis model, which consists of a 1:1 mixture of the Gram-negative bacterium *Escherichia coli* O21:H+ (Ayres et al., 2012) and the Gram-positive bacterium *Staphylococcus aureus* subspecies Rosenbach (ATCC 12600) that was originally isolated from human pleural fluid (Cowan et al., 1954). Fluoxetine pre-treatment protected against infection-induced mortality in a dose-dependent manner (**Figure 1B**). In addition to promoting survival, fluoxetine pre-treatment protected from clinical signs of disease including morbidity and hypothermia (**Figure 1C-D**). Given that 40 mg/kg pretreatment conferred the highest level of protection, we used this dose for our studies.

**Figure 1:**
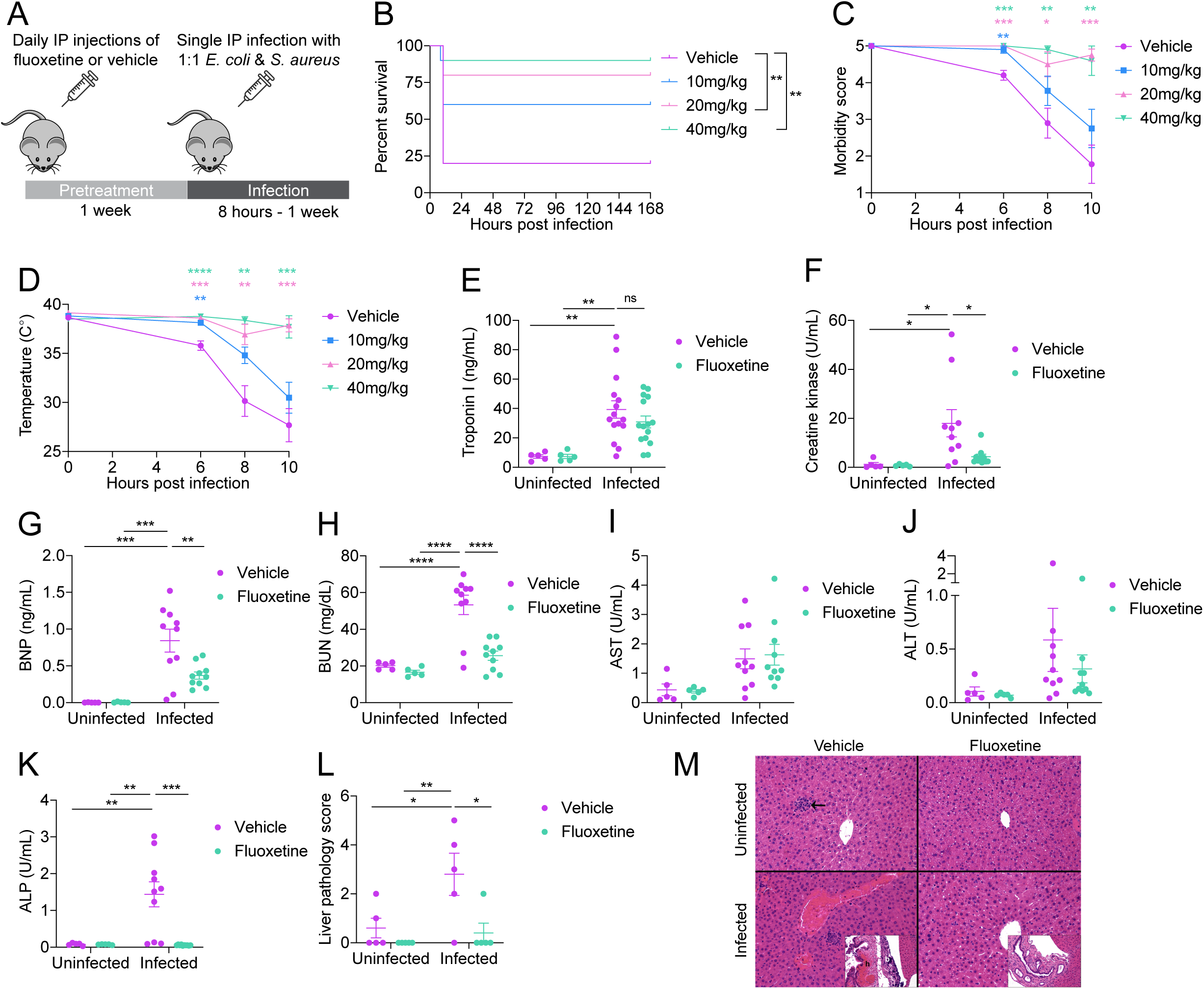
Fluoxetine protects from sepsis-induced disease and mortality. (A) Schematic of experimental approach. C57BL/6J mice were treated for one week with daily intraperitoneal vehicle or fluoxetine injections then intraperitoneally infected with a 1:1 mixture of *E. coli* and *S. aureus*. (B-D) (B) Survival, (C) morbidity, and (D) temperature of vehicle and fluoxetine treated mice infected with polymicrobial sepsis. n=10 per condition, one independent experiment. For survival, log-rank analysis. For morbidity and temperature, two-way ANOVA with Dunnett’s multiple comparisons test. (E-K) Circulating organ damage markers of vehicle and fluoxetine treated mice infected with polymicrobial sepsis. Infected samples collected 8-10 hours post infection. n=5-15 per condition, two-three independent experiments combined. (E) Troponin I. (F) Creatine kinase. (G) Brain natriuretic peptide. (H) Blood urea nitrogen. (I) Aspartate transaminase. (J) Alanine transaminase. (K) Alkaline phosphatase. Two-way ANOVA with Tukey’s multiple comparisons test. (L) Liver pathology of vehicle and fluoxetine treated mice infected with polymicrobial sepsis. Infected samples collected 10 hours post infection. Liver pathology score is combined necrosis, hemorrhage, and congestion scores. Arrow shows a microgranuloma/microabscess, which were seen occasionally in mice from all groups and represent an incidental background lesion. Inset of the worst infected vehicle animal shows acute hemorrhage (h) and bacteria (b) in the region surrounding the gall bladder, compared to the worst fluoxetine infected animal showing no hemorrhage or bacteria. n=5 per condition, one independent experiment. There is mild microvesicular cytoplasmic vacuolation in the fluoxetine treated mice. Two-way ANOVA with Tukey’s multiple comparisons test. In all panels data represent mean ± SEM. * p<0.05, ** p<0.01, *** p<0.001, **** p< 0.0001.

Sepsis is frequently associated with the development of multiorgan dysfunction and failure (Gotts and Matthay, 2016; Hotchkiss et al., 2016). We performed a clinical pathology analysis to understand whether fluoxetine protection from sickness and mortality was associated with less severe organ damage. We found that both vehicle treated and fluoxetine treated infected mice exhibited a comparable elevation in circulating levels of Troponin I, which is a marker of cardiac muscle damage (**Figure 1E**). Fluoxetine treated infected mice exhibited reduced circulating levels of other markers of heart damage compared to vehicle treated infected mice including Creatine Kinase (CK), (which is a marker of both cardiac and skeletal muscle damage) (**Figure 1F**), as well as Brain Natriuretic Peptide (BNP), which is released in response to ventricular wall stretch and during heart failure (Mukoyama et al., 1990) (**Figure 1G**). Together these results demonstrate that while fluoxetine and vehicle-treated mice appear to exhibit similar levels of Troponin I, fluoxetine pre-treatment protects against ventricular stretch and possibly cardiac failure during sepsis. Additionally, we found that fluoxetine treated infected mice were protected from elevated levels of blood urea nitrogen (BUN), which is a marker of kidney damage (**Figure 1H**). Finally, we found that circulating levels of aspartate transaminase (AST) and alanine transaminase (ALT), which are markers of hepatic damage, were elevated to comparable levels in both vehicle treated and fluoxetine treated mice during infection (**Figure 1I-J**). By contrast, fluoxetine treatment protects from elevating circulating levels of an additional marker of liver (and bone) damage, alkaline phosphatase (ALP) (**Figure 1K**). To better understand whether fluoxetine protects against sepsis induced liver damage, we conducted histopathological analysis on livers at 10 hours post-infection. We found that fluoxetine treated infected mice exhibited less necrosis, congestion, and hemorrhage, suggesting fluoxetine pretreatment does indeed confer protection against sepsis-induced liver damage in our model (**Figure 1L-M** and **Supplemental Figure 1A-C**). Together these data demonstrate that fluoxetine pre-treatment protects from sepsis-induced sickness, multi-organ damage, and death.

### Fluoxetine promotes both pathogen resistance and host-pathogen cooperation

To determine the contribution of fluoxetine treatment to antagonistic and cooperative defenses, we measured the colony forming units (CFUs) of *E. coli* and *S. aureus* in the target organs of the pathogens at 8 hours post-infection. Fluoxetine treated mice had significantly lower total pathogen burdens in their liver, spleen, kidneys, and lungs, but not the heart (**Figure 2A**). *E. coli* and *S. aureus* levels followed similar trends in these organs (**Supplemental Figure 1D-E**). These data indicate that fluoxetine treatment has antimicrobial effects either via direct actions on the pathogen and/or by heightening the host resistance response to the infection. Indeed, in agreement with published studies we found that fluoxetine inhibited *E. coli* and *S. aureus* growth at near-clinically relevant concentrations *in vitro* (**Supplemental Figure 1F-G**) (Batista de Andrade Neto et al., 2019; Bolo et al., 2004), suggesting fluoxetine’s direct antimicrobial effects may contribute to the observed decrease in pathogen burden.

**Figure 2:**
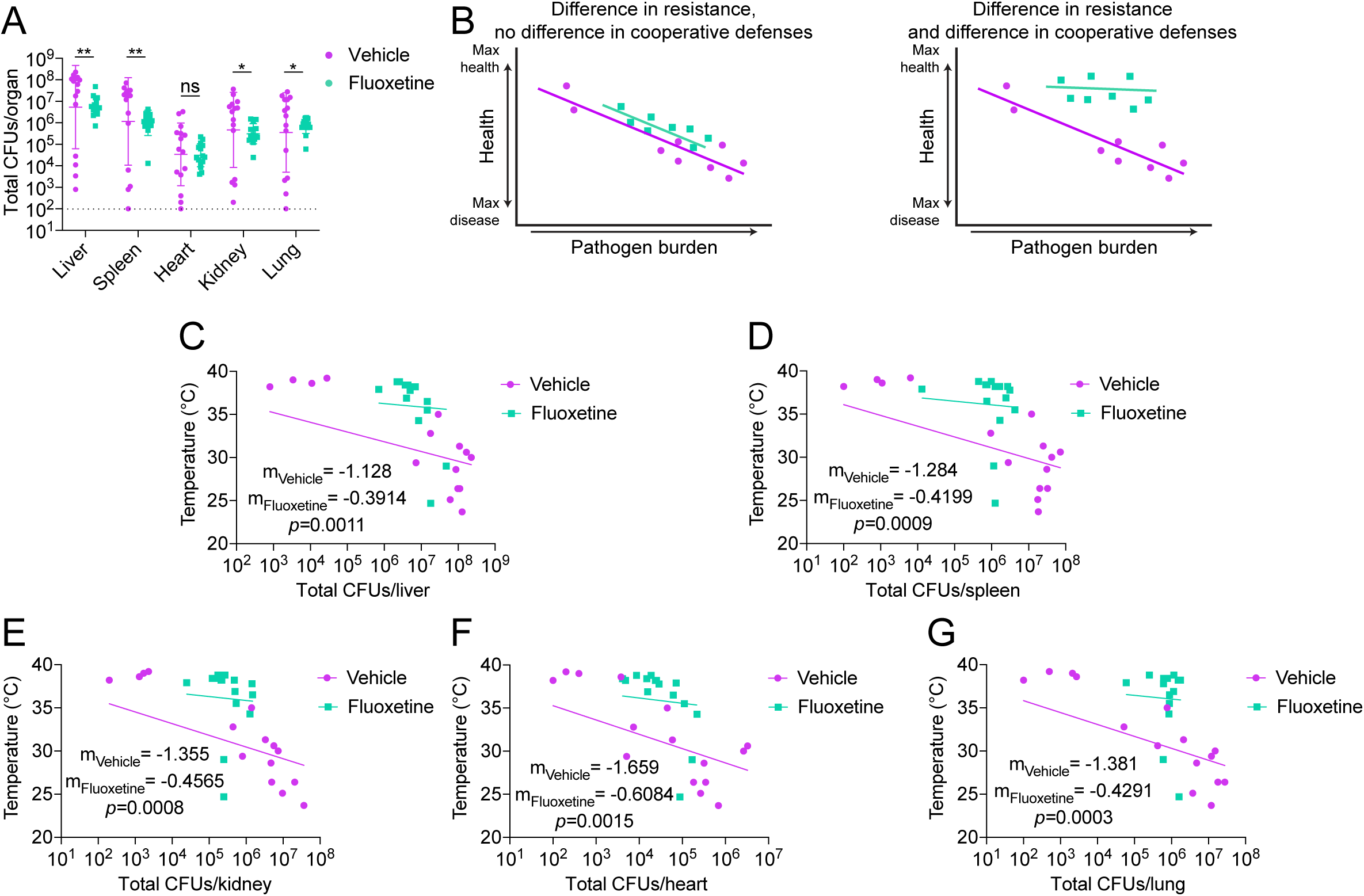
Fluoxetine promotes both pathogen resistance and host-pathogen cooperation. (A) Total pathogen burden analysis from vehicle or fluoxetine treated mice at 8-10 hours post infection. n = 15 per condition, three independent experiments. Data represent geometric mean ± geometric SD. Unpaired t-tests. (B) Hypothetical reaction norms showing differences in resistance alone (left) and resistance and cooperation (right) curves. (C-G) Reaction norm analyses plotting body temperature at the time of dissection against total CFUs for mice in (A). (C) liver (D) spleen (E) Kidney (F) Heart (G) Lung. Semilog linear regression, y-intercept constrained to average of uninfected temperature, Extra sum-of-squares F Test to compare slopes. * p<0.05, ** p<0.01.

Because there were differences in pathogen burdens in almost all the target organs, we needed to employ a reaction norm analysis to determine the contributions of fluoxetine treatment to host-pathogen cooperation (Ayres, 2020a; Råberg et al., 2009). This involves plotting host health against pathogen burdens and examining how host health changes as pathogen burdens change for each treatment group. A shallower slope indicates a better ability to maintain health over a range of pathogen, and therefore improved adaptation to the infected state (**Figure 2B**). We generated reaction norms for each organ using body temperature at 8 hours post-infection as our readout for health and plotted against the pathogen burdens observed for the respective organ at the same time point. We found for all organs that fluoxetine-treated infected mice exhibited shallower slopes compared to vehicle-treated infected mice, in addition to the expected shift along the diagonal that indicates differences in pathogen burdens (**Figure 2C-G** and **Supplemental Figure 1H-Q**). This indicates that fluoxetine treatment facilitates host adaptation to the infected state, and therefore host-pathogen cooperation via an anti-virulence and/or disease tolerance mechanism(s). Taken together, our data suggest that fluoxetine protects from sepsis induced disease and mortality via multiple strategies including antimicrobial effects and also by promoting host-pathogen cooperation.

### Fluoxetine mediated protection from sepsis is serotonin independent

Peripheral serotonin has been reported to contribute to sepsis pathology and mortality in murine models of sepsis (Duerschmied et al., 2013; Zhang et al., 2017). The majority of peripheral serotonin is produced by enterochromaffin cells. In these cells, Tryptophan hydroxylase 1 (TPH1) converts tryptophan to 5-hydroxytryptophan (5-HTP), which is then decarboxylated to generate serotonin (5-HT). Serotonin is then secreted luminally into the gut and basally into circulation (Berger et al., 2009). Once in circulation, serotonin is taken up and sequestered by platelets through the serotonin transporter. Upon activation, platelets release serotonin. SSRIs deplete peripheral serotonin by inhibiting both its export out of enterochromaffin cells and its uptake into platelets (Blardi et al., 2002). We measured serotonin levels in infected fluoxetine treated wild-type mice and compared to *Tph1^-/-^* mice that genetically lack peripheral serotonin. Fluoxetine treated wild-type mice exhibited reduced levels of serotonin in the liver, spleen, lung, and heart comparable to what we observed in organs harvested from *Tph1^-/-^* mice (**Figure 3A-D**). Interestingly, kidney serotonin levels were equivalent across all groups, albeit kidneys had the lowest level of all organs assayed (**Figure 3E**). Furthermore, serum serotonin levels were reduced in fluoxetine treated infected mice to comparable levels as we observed in serum from *Tph1^-/-^* mice (**Figure 3F**). Additionally, at earlier time points throughout polymicrobial sepsis infection, vehicle treated wild-type mice exhibited elevated levels of circulating serotonin, while fluoxetine treated infected wild-type mice were protected from this infection induced increase in circulating serotonin levels (**Supplemental Figure 2A-B**). These differences were independent of platelet activation as measured by thrombocytopenia (**Supplemental Figure 2C-D**). To test the hypothesis that fluoxetine mediated protection from sepsis was dependent on its actions on serotonin, we first measured susceptibility of *Tph1^-/-^* mice challenged with polymicrobial sepsis. We found that *Tph1^-/-^* mice were not protected from disease and mortality compared to *Tph1^+/+^* littermates (**Figure 3G-I**). We then tested whether the protective effects of fluoxetine treatment were abolished in *Tph1^-/-^*mice. Surprisingly, we found that 100% of *Tph1^-/-^* mice treated with fluoxetine were protected from sepsis induced mortality, while ∼50% of vehicle-treated infected *Tph1^-/-^* mice succumbed to the challenge (**Figure 3J**). Fluoxetine treatment also protected *Tph1^-/-^* mice from clinical signs of disease (**Figure 3K-L**). Taken together, these data demonstrate that in mice lacking peripheral serotonin, fluoxetine still confers protection against sepsis induced morbidity and mortality, suggesting the fluoxetine-mediated protection during polymicrobial sepsis is serotonin-independent.

**Figure 3:**
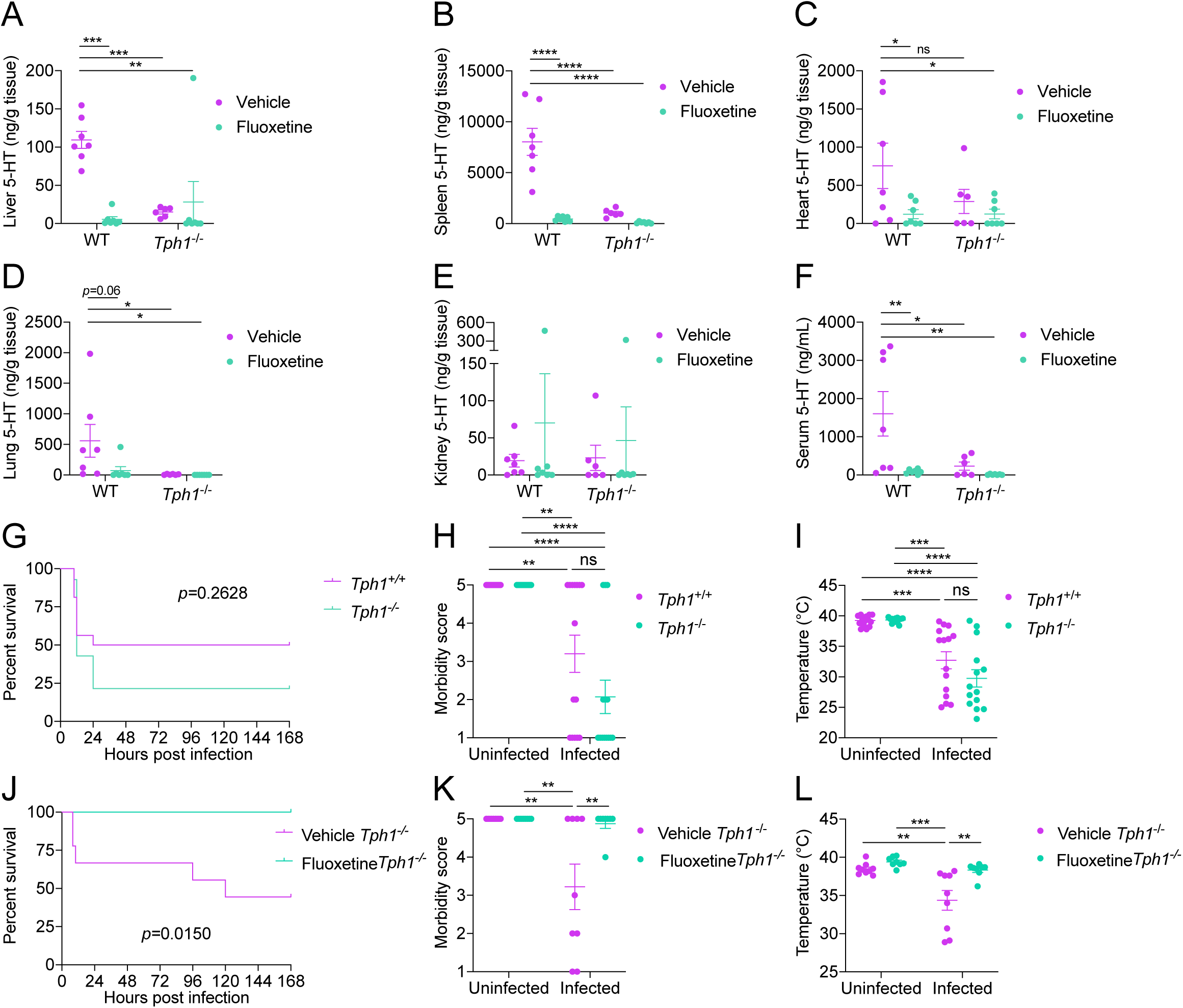
Fluoxetine mediated protection from sepsis is serotonin independent. (A-F) Serotonin (5-HT) levels of wildtype and *Tph1*^-/-^ mice at 8-10 hours post infection in vehicle or fluoxetine treated mice infected with polymicrobial sepsis. (A) Liver (B) Spleen (C) Heart (D) Lung (E) Kidney (F) Serum. n=6-7 per condition, two independent experiments combined. Two-way ANOVA with Dunnett’s multiple comparisons test against WT Vehicle. (G-I) (G) Survival, (H) most severe morbidity score exhibited over course of infection by each mouse, and (I) minimum temperature exhibited over course of infection by each mouse of *Tph1*^+/+^ or *Tph1*^-/-^ littermates infected with polymicrobial sepsis. n=14-16 per condition, four independent experiments combined. For survival, Log-rank analysis. For morbidity score and temperature, Two-way ANOVA with Tukey’s multiple comparisons. (J-L) (J) Survival, (K) most severe morbidity score exhibited over course of infection by each mouse, and (L) minimum temperature exhibited over course of infection by each mouse of *Tph1*^-/-^ mice treated with vehicle or fluoxetine infected with polymicrobial sepsis. n=8-9 per condition, three independent experiments combined. For survival, Log-rank analysis. For morbidity score and temperature, Two-way ANOVA with Tukey’s multiple comparisons. In all panels data represent mean ± SEM. * p<0.05, ** p<0.01, *** p<0.001, **** p<0.0001.

### IL-10 is necessary for fluoxetine mediated protection from sepsis

Sepsis pathogenesis is largely driven by an over-exuberant inflammatory response to infection (Angus and van der Poll, 2013; Gotts and Matthay, 2016; Van Der Poll et al., 2017). *In vitro* and *in vivo* studies have demonstrated that fluoxetine has anti-inflammatory effects (Abdel-Salam et al., 2004; Liu et al., 2011; Maes et al., 2005; Roumestan et al., 2007). In a rat endotoxemia model, fluoxetine treatment protected from elevated circulating levels of the pro-inflammatory cytokines TNFα, IL-6, and IL-1β (Mojiri-Forushani et al., 2023). In our polymicrobial sepsis model, at two hours post-infection, we found that fluoxetine treated mice had equivalent induction of TNFα and IL-6, and significantly higher levels of serum IL-1β compared to vehicle treated infected mice (**Figure 4A-C**). This inflammatory response was resolved in fluoxetine-treated mice by 8 hours post-infection, while vehicle-treated infected mice exhibited dramatically increased levels of these cytokines at this late time point with a similar magnitude and pattern exhibited by human septic patients (Damas et al., 1992) (**Figure 4A-C**). Hepatic transcript levels of these cytokines showed similar induction and kinetic patterns as we observed in serum (**Supplemental Figure 2E-G**).

**Figure 4:**
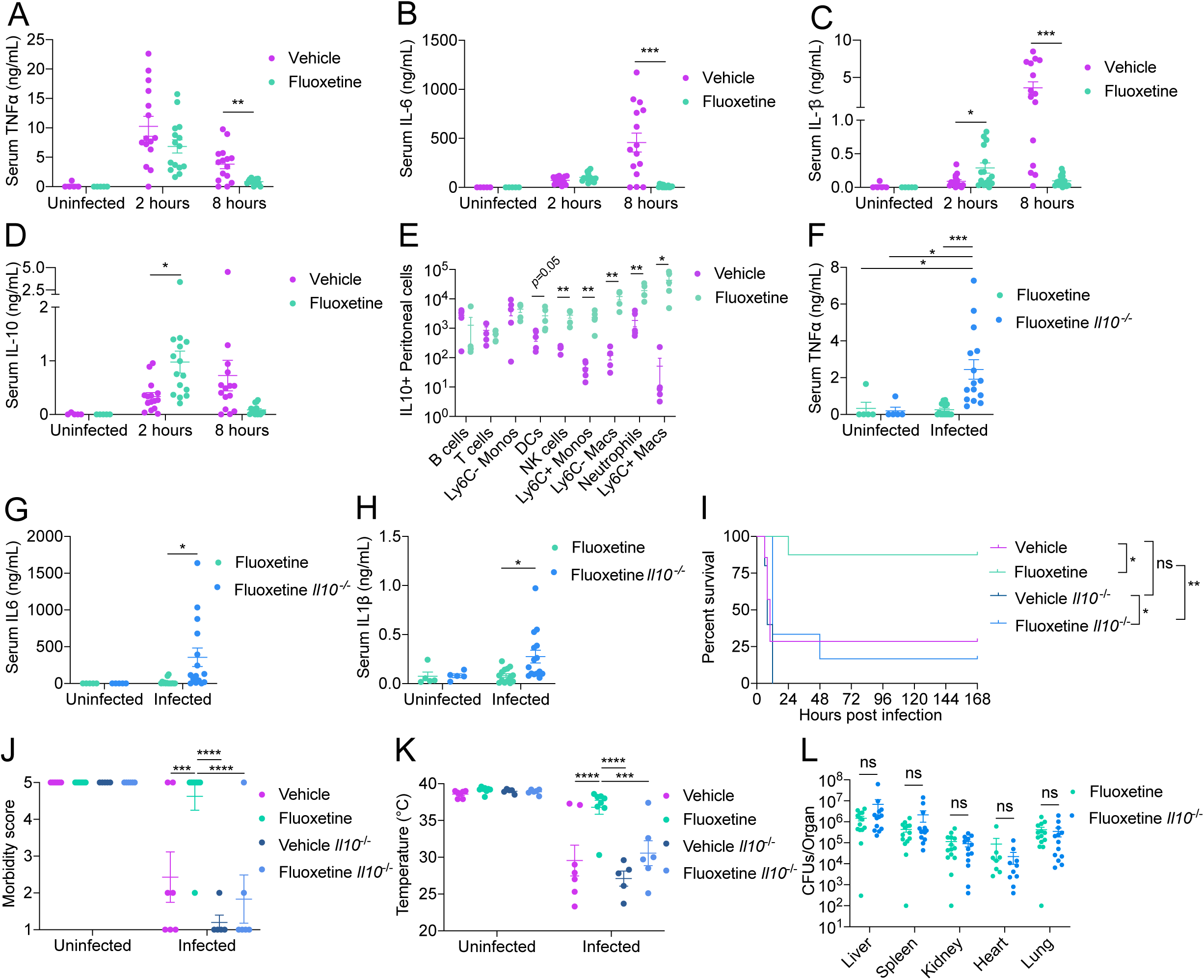
IL-10 is necessary for fluoxetine mediated protection from sepsis. (A-D) Circulating levels of (A) TNFα, (B) IL-6, (C) IL-1β, and (D) IL-10 in vehicle or fluoxetine treated mice infected with polymicrobial sepsis. n=5-15 per condition, three independent experiments combined. Data represent mean ± SEM. Unpaired t-tests with Holm-Sidak multiple comparisons correction. (E) Flow cytometry analysis of peritoneal lavage cells 2 hours post infection from vehicle or fluoxetine treated mice infected with polymicrobial sepsis. n=5 per condition, one independent experiment. Data represent mean ± SEM. Unpaired t-tests. (F-H) Circulating levels of (F) TNFα, (G) IL-6, and (H) IL-1β at 10 hours post infection in wildtype or *Il10^-/-^* mice pre-treated with fluoxetine infected with polymicrobial sepsis. n=5-11 per condition, two independent experiments combined. Data represent mean ± SEM. Two-way ANOVA with Tukey’s multiple comparisons test. (I-K) (I) Survival, (J) most severe morbidity score exhibited over course of infection by each mouse, and (K) minimum temperature exhibited over course of infection by each mouse of wildtype or *Il10^-/-^* mice treated with vehicle or fluoxetine infected with polymicrobial sepsis. n = 5-8 per condition, two independent experiments. Data represent mean ± SEM. For survival, Log-rank analysis. For morbidity and temperature, Two-way ANOVA with Tukey’s multiple comparison test. (J) Total pathogen burden analysis at 10 hours post infection from wildtype or *Il10^-/-^* mice treated with fluoxetine infected with polymicrobial sepsis 10 hours post infection. n=8-14 per condition, two or three independent experiments combined. Data represent geometric mean ± geometric SD. Unpaired t-tests. * p<0.05, ** p<0.01, *** p<0.001, **** p<0.0001.

Pro-inflammatory cytokine production can be limited by concurrent anti-inflammatory cytokine production (Van Der Poll et al., 2017). At 2 hours post infection, fluoxetine pre-treated mice exhibited significantly greater induction of circulating and tissue levels of IL-10 compared to vehicle-treated mice (**Figure 4D** and **Supplemental Figure 2H**). Furthermore, we found greater peritoneal recruitment of multiple IL-10+ innate immune cell populations, including NK cells, neutrophils, Ly6C+ monocytes, Ly6C-macrophages, and Ly6C+ macrophages, from fluoxetine treated infected mice 2 hours post-infection (**Figure 4E** and **Supplemental Figure 2I-K**). We did not observe significantly different percentages of IL-10+ cells in these populations (**Supplemental Figure 2I**). We also did not observe changes in numbers or percentage of IL-10+ Ly6C-monocytes, T cells, and B cells (**Figure 4E** and **Supplemental Figure 2I-K**). To test the importance of IL-10 for fluoxetine mediated protection from sepsis, we subjected *Il10^-/-^* mice to our fluoxetine treatment paradigm (**Figure 1A**) and measured pro-inflammatory cytokine production. The anti-inflammatory effects of fluoxetine at the later stages of infection were abrogated in *Il10^-/-^* mice (**Figure 4F-H** and **Supplemental Figure 2L-N**). Furthermore, fluoxetine pre-treatment did not protect *Il10^-/-^* mice from sepsis induced morbidity and mortality (**Figure 4I-K**). Finally, from our pathogen burden analysis, we found that there were no differences in total pathogen burden, nor independently *E. coli* or *S. aureus* burden, between fluoxetine treated infected wild type and *Il10^-/-^* mice (**Figure 4L** and **Supplemental Figure 3A-B**). Instead, as revealed from our reaction norm analysis, fluoxetine treated *Il10^-/-^* mice exhibited steeper slopes both when analyzing total CFUs or either *E. coli* or *S. aureus* CFUs individually (**Supplemental Figure 3C-G**). Together, these data demonstrate that IL-10 is necessary for the anti-inflammatory effects of fluoxetine as well as the protection from sepsis induced disease and mortality mediated by fluoxetine treatment. Finally, our data demonstrate that the ability of fluoxetine to promote host adaptation to the infection and cooperative defenses is dependent on IL-10.

### Fluoxetine protects from sepsis induced hypertriglyceridemia

Sepsis causes disturbances in lipid homeostasis including elevated levels of circulating triglycerides (Cetinkaya et al., 2014). Fluoxetine treatment in patients has been associated with changes in circulating triglycerides and other lipids (Pan et al., 2018). *In vitro*, fluoxetine treatment has been shown to regulate aspects of lipid homeostasis (Xiong et al., 2014). We found that wild-type mice challenged with our polymicrobial sepsis infection exhibited an increase in circulating triglycerides, while fluoxetine mice maintain uninfected levels during infection (**Figure 5A**). Triglyceride levels are regulated exogenously by diet and by endogenous mechanisms. As part of our infection protocol, we withhold food for the first 12 hours during polymicrobial sepsis, thus differences in food consumption cannot contribute to differences in triglyceride metabolism. This suggests that endogenous processes are responsible for the observed differences in triglyceride levels between treatment groups. We first tested whether fluoxetine treatment affected hepatic triglyceride export during infection. To test this, we injected vehicle and fluoxetine-treated mice with Pluronic F-127 during infection and measured serum triglyceride levels every hour post-infection for four hours. Pluronic F-127 inhibits the activity of lipoprotein lipase (LPL), and so any changes detected in serum triglyceride levels is suggested to reflect hepatic triglyceride export (Johnston and Palmer, 1993). Fluoxetine treated infected mice showed a slight but insignificant reduction in triglyceride levels at four hours post-Pluronic F-127 injection (**Figure 5B**). That said, blocking hepatic triglyceride export with the microsomal triglyceride transport protein (Mttp) inhibitor lomitapide did not significantly influence polymicrobial sepsis survival (**Supplemental Figure 4A**). Furthermore, fluoxetine treated mice exhibited no difference in hepatic transcript levels of fatty acid transport (*Cd36)*, lipogenesis (*Fasn, Acaca, Srebp1*), nor triglyceride export related genes (*Apob, Mttp)* (**Figure 5C**).

**Figure 5:**
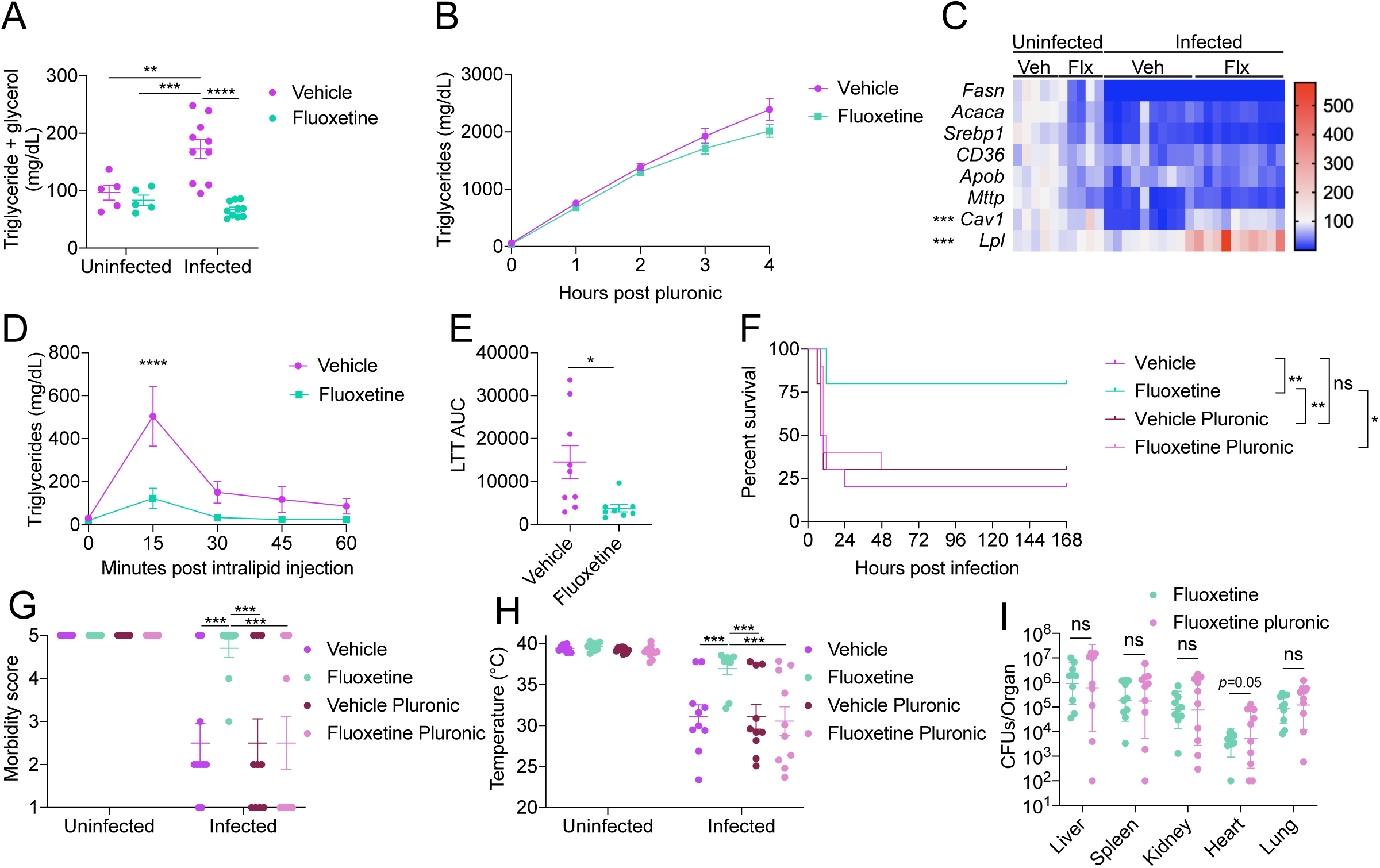
Fluoxetine protects from sepsis induced dyslipidemia. (A) Circulating levels of triglycerides + glycerol at 8-10 hours post infection in vehicle or fluoxetine treated mice infected with polymicrobial sepsis. n=5-10 per condition, two independent experiments combined. Data represent mean ± SEM. Two-way ANOVA with Tukey’s multiple comparisons. (B) Hepatic triglyceride export using the poloxamer method measured starting at 2 hours post infection in vehicle or fluoxetine treated mice infected with polymicrobial sepsis. n=10 per condition, one independent experiment. Data represent mean ± SEM. Unpaired t-tests with Holm-Sidak multiple comparisons correction. (C) Hepatic transcript levels of lipid metabolism-related genes at 8-10 hours post infection in vehicle or fluoxetine treated mice infected with polymicrobial sepsis. n=5-10 per condition, two independent experiments combined. Two-way ANOVA with Tukey’s multiple comparisons, significance stars indicate comparison between infected vehicle and infected fluoxetine. (D-E) Lipid tolerance test (LTT) at 7 hours post infection in vehicle or fluoxetine treated mice infected with polymicrobial sepsis. n=8-9 per condition, two independent experiments combined. (D) serum triglyceride levels and (E) area under the curve analysis of (D). Data represent mean ± SEM. For LTT, two-way ANOVA with Sidak multiple comparisons between vehicle and fluoxetine at each timepoint. For AUC, unpaired student’s t-test. Three of the fluoxetine mice are also plotted in Panels 6E-F. The Flx *Il10*^-/-^ mice in Panels 6E-F were assayed in parallel to one of the Vehicle vs. Fluoxetine repeats in Panels 5D-E. (F-H) (F) Survival, (G) most severe morbidity score exhibited over course of infection by each mouse, and (H) minimum temperature exhibited over course of infection by each mouse of vehicle or fluoxetine treated mice with or without Pluronic F-127 injection at the time of infection with polymicrobial sepsis. n=10 per condition, two independent experiments combined. Data represent mean ± SEM. For survival, log-rank analysis. For morbidity and temperature two-way ANOVA with Tukey’s multiple comparisons. (I) Total pathogen burden analysis at 8-10 hours post infection in fluoxetine treated mice with or without Pluronic F-127 injection at the time of infection with polymicrobial sepsis. n=10 per condition, two independent experiments combined. Data represent geometric mean ± geometric SD. Unpaired t-tests. * p<0.05, ** p<0.01, *** p<0.001, **** p< 0.0001.

Increased peripheral uptake of triglycerides is an additional endogenous process that will regulate circulating triglyceride levels. To test whether fluoxetine regulates triglyceride uptake during infection we performed a lipid tolerance test (LTT) by injecting infected mice intravenously with a bolus of intralipid and measuring the clearance of triglycerides from the blood. We found that infected mice that received fluoxetine treatment were more lipid tolerant compared to vehicle treated mice, demonstrating that fluoxetine increases peripheral uptake of triglycerides (**Figure 5D-E**). In corroboration with this result, fluoxetine-treated mice exhibited increased hepatic transcript levels of *Lpl* and *Cav1*, genes responsible for endocytosis of cholesterol and triglycerides (**Figure 5C**). To determine whether triglyceride uptake is necessary for fluoxetine mediated protection, we administered Pluronic F-127 to mice during infection to inhibit LPL activity and triglyceride uptake, and monitored disease progression. We found that Pluronic F-127 treatment abolished the protective effects of fluoxetine in polymicrobial sepsis challenged animals (**Figure 5F**). Fluoxetine/pluronic treated infected mice exhibited increased mortality and more severe clinical signs of disease compared to fluoxetine treated infected mice (**Figure 5G-H**). We found that there were no differences in total pathogen burden between fluoxetine and fluoxetine/pluronic mice in liver, spleen, kidney, and lung, and there was a near significant increase in the hearts of fluoxetine/pluronic mice (**Figure 5I**). When we analyzed each pathogen independently, we found that there was no difference in *S. aureus* CFUs between conditions while there was an increase in *E. coli* CFUs in the livers, kidneys, hearts, and lungs of fluoxetine/pluronic mice (**Supplemental Figure 4B-C**). Our reaction norm analysis demonstrates that fluoxetine/pluronic mice exhibit steeper slopes across all five organs assayed both when analyzing total CFUs and either *E. coli* or *S. aureus* CFUs individually (**Supplemental Figure 4D-H**). Together these data demonstrate that fluoxetine promotes triglyceride uptake during polymicrobial sepsis, which is necessary to promote host-pathogen cooperation, resulting in protection from infection induced morbidity and mortality.

### IL-10 is necessary for fluoxetine mediated protection from hypertriglyceridemia during sepsis

We next asked what the relationship is between the fluoxetine mediated effects on IL-10 and triglycerides. We first tested whether triglyceride uptake is required for fluoxetine-induced IL-10 production. Pluronic decreased the number of IL-10+ peritoneal monocytes and decreased the percentage of IL-10+ peritoneal monocytes and dendritic cells 2 hours post infection in fluoxetine treated mice, but did not impact recruitment of IL-10+ cells or IL-10 production in any other cell type assayed (**Figure 6A** and **Supplemental Figure 4I**). Conversely, pluronic did not impact circulating serum levels of IL-10, or hepatic *Il10* transcription 2 hours post infection in fluoxetine treated mice (**Figure 6B-C**). Together these data indicate that some peritoneal cell types require triglyceride uptake to increase IL-10 production during sepsis in fluoxetine treated mice, but ultimately the fluoxetine-mediated increase in circulating and hepatic IL-10 during sepsis is independent of triglyceride uptake. We next tested whether IL-10 production is required for maintenance of triglyceride homeostasis during sepsis. While we found no differences in the levels of circulating triglycerides when uninfected, fluoxetine treated infected *Il10^-/-^* mice exhibited significantly elevated levels of circulating triglycerides compared to fluoxetine treated infected wild-type mice (**Figure 6D**). From our LTT analysis, we found that fluoxetine treated *Il10^-/-^* infected mice were less lipid tolerant compared to fluoxetine treated infected wild-type mice (**Figure 6E-F**) demonstrating that IL-10 is necessary for fluoxetine mediated increased peripheral uptake of lipids during infection. Taken together, our data suggest that fluoxetine mediated effects on lipid uptake and protection from hypertriglyceridemia are dependent on IL-10. By contrast, increased lipid uptake and protection from hypertriglyceridemia is not necessary for the fluoxetine mediated effects on IL-10 during infection.

**Figure 6:**
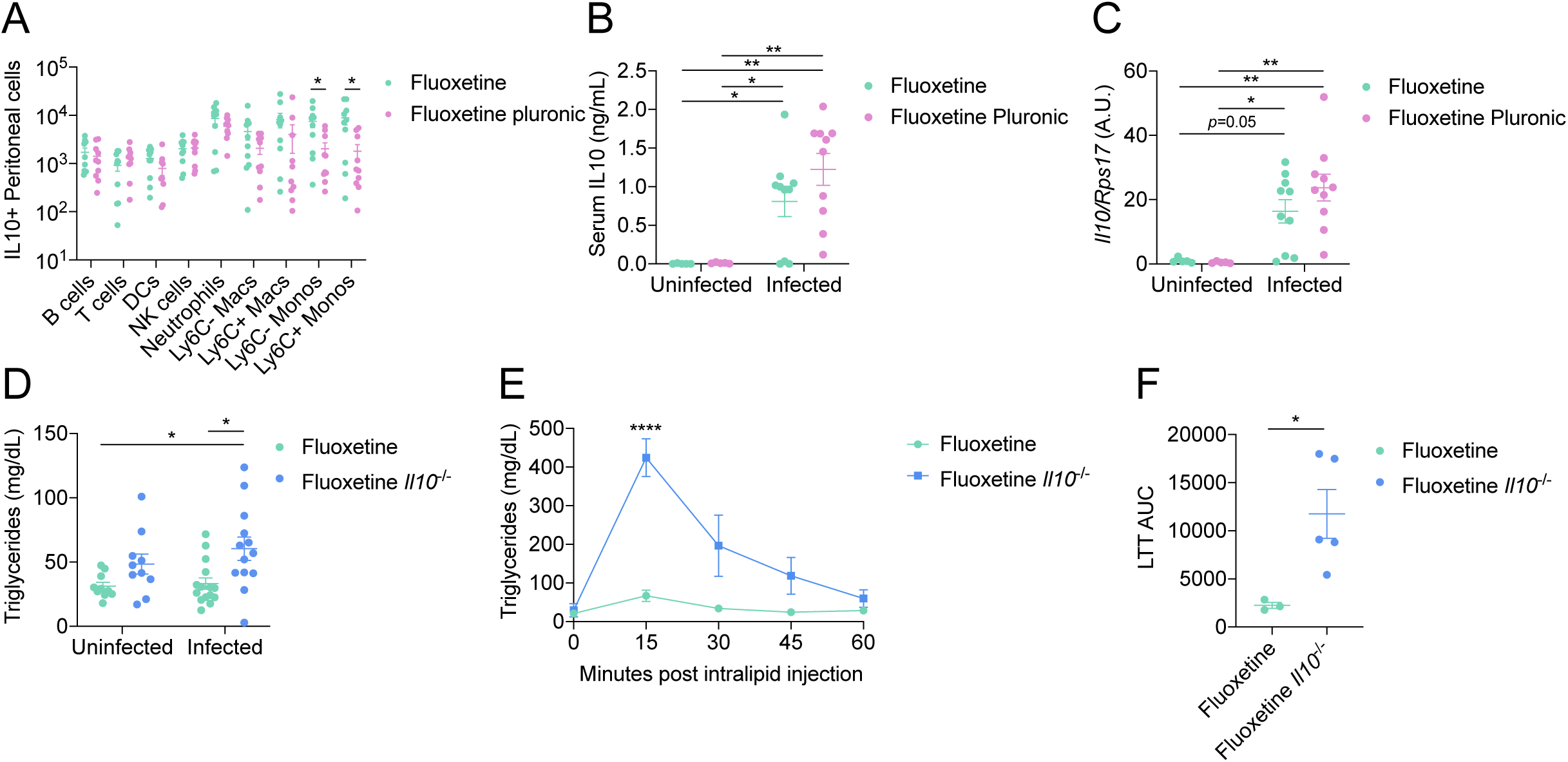
IL-10 is necessary for fluoxetine mediated protection from hypertriglyceridemia during sepsis. (A-C) (A) Peritoneal lavage flow cytometry, (B) circulating IL-10, and (C) hepatic Il10 transcription at 2 hours post infection in fluoxetine treated mice injected with Pluronic F-127 at the time of infection with polymicrobial sepsis. n=5-10 per condition, one or two independent experiments combined. For flow cytometry, unpaired t-tests. For circulating IL-10 and hepatic Il10, two-way ANOVA with Tukey’s multiple comparisons. Circulating levels of triglycerides at 8-10 hours post infection in fluoxetine treated WT or *Il10^-/-^* mice infected with polymicrobial sepsis. n=10-15 per condition, three independent experiments combined. Two-way ANOVA with Tukey’s multiple comparisons. (E-F) Lipid tolerance test (LTT) at 7 hours post infection in fluoxetine treated WT or *Il10^-/-^* mice infected with polymicrobial sepsis. n=3-5 per condition, one independent experiment. Three of the fluoxetine mice are also plotted in Panels 5D-E. The Flx *Il10*^-/-^ mice in Panels 6E-F were assayed in parallel to one of the Vehicle vs. Fluoxetine repeats in Panels 5D-E. For LTT, two-way ANOVA with Sidak multiple comparisons between vehicle and fluoxetine at each timepoint. For AUC, unpaired student’s t-test. In all panels data represent mean ± SEM. * p<0.05, ** p<0.01, **** p< 0.0001.

### IL-10 and protection from hypertriglyceridemia are necessary for fluoxetine mediated metabolic changes in the septic heart

To begin to understand how fluoxetine protects from sepsis induced disease and mortality, we initiated lines of investigation to understand how fluoxetine regulation of IL-10 and triglycerides protect from organ dysfunction and damage. We chose to focus our efforts on the heart because we found that fluoxetine treated mice were protected from elevated levels of BNP, suggesting protection from ventricular stretch and possibly heart failure (**Figure 1G**). Furthermore, previous research has demonstrated a cardioprotective effect of SSRIs (Tagashira et al., 2010). Cardiac metabolism changes dramatically in the septic heart and dysregulation in cardiac metabolism is proposed to be a driver of sepsis mortality in humans (Rudiger and Singer, 2007). Under physiological conditions, fatty acids (FAs) are the main energy substrate for the heart. During conditions of heart failure, there is a reduction in fatty acid oxidation and an increase in use of alternative energy sources (Doenst et al., 2013). To determine how fluoxetine regulates fatty acid oxidation (FAO) in the septic heart, we first analyzed the levels of active Acetyl-CoA Carboxylase 1/2 (ACC1/2), a critical regulator of FAO. Activity of ACC1/2 is regulated by its phosphorylation state: dephosphorylation activates the enzyme and phosphorylation inactivates the enzyme. We found no difference in the ratio of pACC1/2:total ACC1/2 across any treatment groups (**Figure 7A** and **Supplemental Figure 5A**). We also found no difference in the expression of genes involved in fatty acid oxidation (FAO) including *Pgc1a, Pgc1b, Acadm, Erra,* and *Cpt1b* in the hearts of fluoxetine treated and vehicle treated infected mice during infection (**Figure 7B**). Taken together, these data suggest there are no differences in the levels of FAO in the two different treatment groups during sepsis.

**Figure 7:**
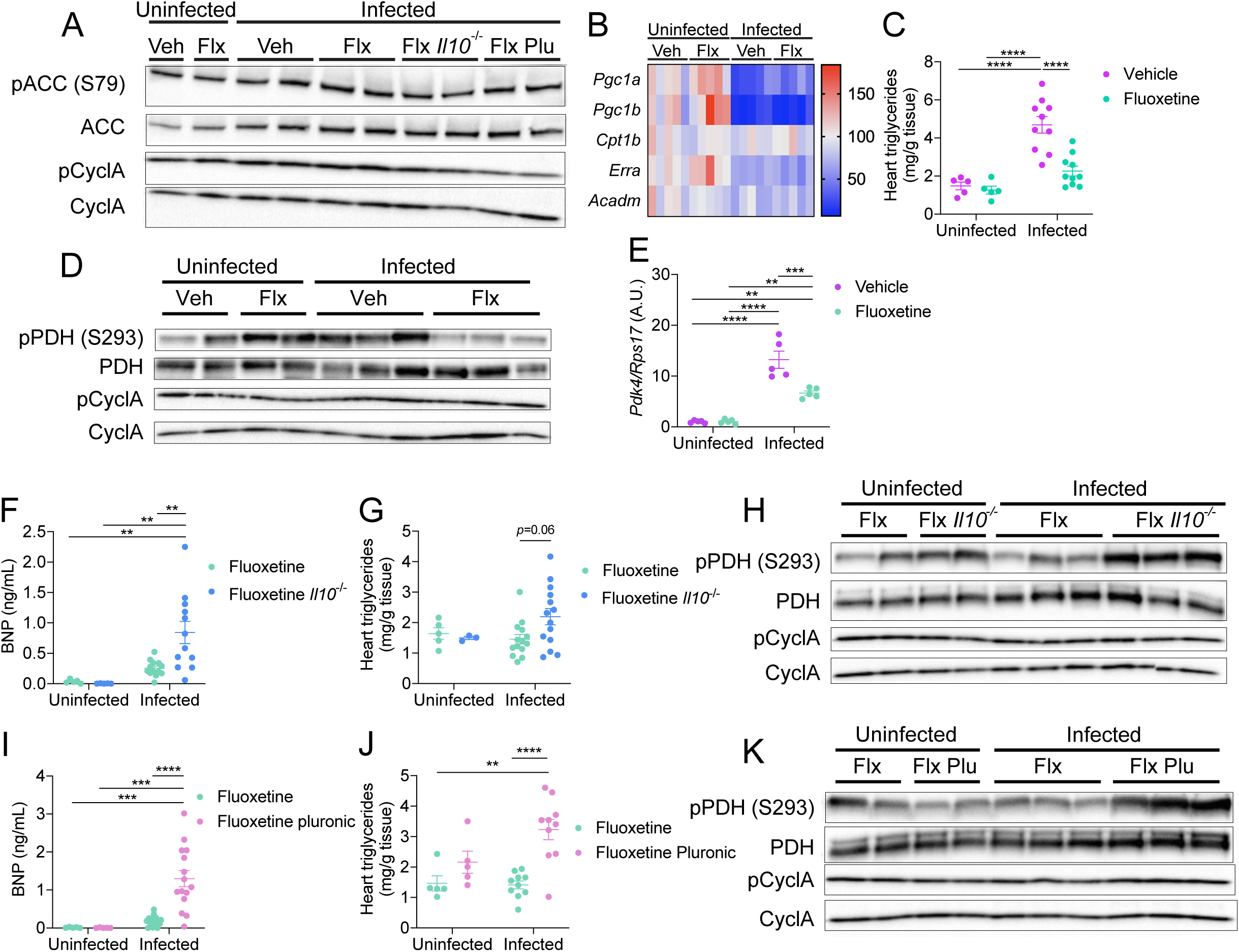
IL-10 and protection from hypertriglyceridemia are necessary for fluoxetine mediated metabolic changes in the septic heart. (A) Western blot of acetyl-coa carboxylase (ACC), phospho-ACC, and cyclophilin A of hearts at 8-10 hours post infection in vehicle or fluoxetine treated WT or *Il10*^-/-^ mice injected with Pluronic F-127 at the time of infection with polymicrobial sepsis. Blot labeled “tCyclA” is the loading control for the total ACC blot. Blot labeled “pCyclA” is the loading control for the pACC blot. n=1-2 per condition, representative of three independent experiments. Uncropped blots are shown in in Supplemental Figure 5. (B) Cardiac fatty acid oxidation-related transcript levels at 10 hours post infection in vehicle or fluoxetine treated mice infected with polymicrobial sepsis. n=5 per condition, one independent experiment. Two-way ANOVA with Tukey’s multiple comparisons, no significant results comparing infected vehicle to infected fluoxetine. (C) Cardiac triglyceride levels at 8-10 hours post infection in vehicle or fluoxetine treated mice infected with polymicrobial sepsis. n=5-10 per condition, two independent experiments combined. Two-way ANOVA with Tukey’s multiple comparisons. (D) Western blot of pyruvate dehydrogenase complex (PDH), phospho-PDH, and cyclophilin A of hearts at 8-10 hours post infection in vehicle or fluoxetine treated mice infected with polymicrobial sepsis. Blot labeled “CyclA” is the loading control for total PDH blot. Blot labeled “pCyclA” is the loading control for the pPDH blot. n=2-3 per condition, representative of three independent experiments. Uncropped blots are shown in in Supplemental Figure 5. (E) Cardiac Pdk4 transcript levels at 10 hours post infection in vehicle or fluoxetine treated mice infected with polymicrobial sepsis. Same mice as in (B). Two-way ANOVA with Tukey’s multiple comparisons. (F-H) (F) Circulating BNP, (G) cardiac triglycerides, and (H) western blot of PDH, phospho-PDH, and cyclophilin A of hearts at 8-10 hours post infection in fluoxetine treated wildtype or *Il10^-/-^* mice infected with polymicrobial sepsis. Serum BNP n=5-15 per condition, three independent experiments combined. Two-way ANOVA with Tukey’s multiple comparisons. Cardiac triglycerides n=3-14 per condition, three independent experiments combined. Two-way ANOVA with Tukey’s multiple comparisons. Western blots n=2-3 per condition, representative of two independent experiments. Original blots are shown in Supplemental Figure 5. (I-K) (I) Circulating BNP, (J) cardiac triglycerides, and (K) western blot of PDH, phospho-PDH, and cyclophilin A of hearts at 8-10 hours post infection in fluoxetine treated mice injected with water or Pluronic F-127 at the time of infection with polymicrobial sepsis. Serum BNP n=5-15 per condition, three independent experiments combined. Two-way ANOVA with Tukey’s multiple comparisons. Cardiac triglycerides n=5-10 per condition, two independent experiments combined. Two-way ANOVA with Tukey’s multiple comparisons. Western blots n=2-3 per condition, representative of two independent experiments. Original blots are shown in in Supplemental Figure 5. In all panels data represent mean ± SEM. * p<0.05, ** p<0.01, *** p<0.001, **** p< 0.0001.

Elevated circulating triglyceride levels can lead to ectopic accumulation of triglycerides and lipotoxicity in organs including the heart, which has been proposed to perturb cardiac metabolism (Wende and Abel, 2010). Consistent with this, we found that vehicle treated infected mice exhibited triglyceride accumulation in the heart, while fluoxetine treatment protected mice from cardiac triglyceride accumulation during infection (**Figure 7C**). In heart failure and the septic heart, there is an increase in the rate of glucose uptake and glycolysis, but typically not an increase in the rate of glucose oxidation (Zheng et al., 2017). Shifting to increased glucose oxidation is proposed to be of benefit to the septic heart because glucose is a more efficient energy source in terms of oxygen consumption compared to FAs (Doenst et al., 2013). We found that vehicle treated infected mice had an increase in the ratio of pPDH:total PDH compared to vehicle treated uninfected wild-type mice, indicating reduced levels of pyruvate fueling the TCA cycle and subsequent oxidative phosphorylation (**Figure 7D** and **Supplemental 5B**). By contrast, fluoxetine treatment protected against inhibition of PDH activity in the heart during sepsis (**Figure 7D** and **Supplemental 5C**). Consistent with this, *Pdk4,* the major kinase responsible for cardiac PDH phosphorylation, was elevated in vehicle-treated mice during infection compared to fluoxetine-treated mice (**Figure 7E**). Importantly, the effects of fluoxetine treatment on cardiac failure, triglyceride accumulation, and pyruvate oxidation were dependent on both IL-10 (**Figure 7F-H** and **Supplemental Figure 5D**) and Lpl activity (**Figure 7I-K** and **Supplemental Figure 5E**). Taken together, our data suggest that fluoxetine promotes metabolic reprogramming in the septic heart and protects from ectopic lipid accumulation, and this is dependent on IL-10 protection against hypertriglyceridemia.

## Discussion

Sepsis is defined as a dysregulated systemic inflammatory response that can lead to multi-organ failure and death (Hotchkiss et al., 2016). Historically, basic sepsis research has focused on characterizing the inflammatory factors driving disease progression, e.g. IL-1β, TNFα, TLR4, etc., with the assumption that blocking these mechanisms of disease in the clinic with either small molecules or monoclonal antibodies will be sufficient to promote survival. Unfortunately, clinical trials based on this approach have been largely unsuccessful. A new perspective for treating sepsis is needed. In addition to characterizing mechanisms of disease pathogenesis, it is critical to identify mechanisms to promote host-pathogen cooperation and develop therapeutic strategies that target these host encoded responses. In the current study, we identified the SSRI fluoxetine as a prophylactic agent that promotes survival in a murine model of polymicrobial sepsis, and revealed the immunometabolic mechanisms by which this occurs.

Fluoxetine has been previously used as a pharmacological tool to study the role of serotonin during sepsis (Duerschmied et al., 2013). However, we found that fluoxetine-mediated protection is independent of serotonin in our sepsis model. Additionally, multiple studies have used orthogonal approaches to study the role of serotonin in sepsis such as *Tph1^-/-^*mice (Duerschmied et al., 2013; Zhang et al., 2017), but we also found that these mice are not protected from mortality and death in our model. There are several factors that may explain the discrepancies between our work and these previously published studies. First, previous studies used LPS and cecal ligation and puncture (CLP) models of sepsis, while we used our polymicrobial sepsis model.

Second, previous studies used male mice while we used females for our study. Notably, while work by Rosen et al. found that fluvoxamine, another SSRI, protects against mortality following LPS or fecal slurry injection by acting through the sigma-1 receptor; the authors did not test the role of serotonin in their model (Rosen et al., 2019). Future work is needed to determine in greater detail under which circumstances serotonin is pathogenic, and what molecular targets SSRIs act through to mediate their protective effects.

Previous reports have suggested that fluoxetine has anti-inflammatory effects (Durairaj et al., 2015; Tynan et al., 2012). In the current study, we found that fluoxetine does not completely abolish the pro-inflammatory response. Instead, fluoxetine controls the degree and duration of the pro-inflammatory response. In fluoxetine-treated mice, we found that at the early stages of infection, there is a comparable to higher level of induction of pro-inflammatory cytokines as vehicle-treated mice. In parallel, fluoxetine-treated mice exhibit a greater induction of the anti-inflammatory cytokine IL-10. However, by the later stages of infection, the inflammatory response has returned to baseline in fluoxetine-treated mice, while it persists in vehicle-treated mice. While we do not know the cell types that are necessary and sufficient for the increase in circulating IL-10, we found that fluoxetine induced greater peritoneal recruitment of several different IL-10+ innate cell types. Future studies are needed to determine the mechanism by which fluoxetine regulates IL-10, and determine the contribution of each of these cell types in fluoxetine’s anti-inflammatory effects on regulating lipid homeostasis and sepsis disease progression.

We found that fluoxetine increases peripheral lipid uptake and protects from hypertriglyceridemia in an IL-10 dependent manner. Previous work has demonstrated that fluoxetine can drive hepatic triglyceride accumulation in non-sepsis contexts (Feng et al., 2012). In agreement with this, we found that there is increased expression of genes involved in lipid uptake including *Lpl* and *Cav1*, in the liver of fluoxetine-treated septic mice, suggesting that fluoxetine and IL-10 may promote normolipidemia by increasing hepatic triglyceride uptake. Increased circulating triglyceride levels can lead to ectopic accumulation of lipids in the heart and development of cardiovascular disease (Miller et al., 2011). Consistent with this, we found that the elevated circulating triglycerides found in vehicle-treated septic mice was associated with cardiac triglyceride accumulation. Lipid accumulation in the heart disrupts the metabolic flexibility of the heart (Drosatos and Schulze, 2013). Indeed, based on indirect measurements, we found that vehicle-treated infected mice had reduced levels of pyruvate fueling the TCA cycle and subsequent oxidative phosphorylation in the heart, while fluoxetine treatment protected against inhibition of PDH activity in the heart during sepsis. We propose that this shift is necessary for protection from septic-induced ventricular stretch and possibly cardiac failure, and contributes to fluoxetine’s protective effects on sepsis induced morbidity and mortality. This is in line with previous work that has demonstrated fluoxetine protects against cardiac damage following aortic constriction (Tagashira et al., 2010).

A previous study using a mouse model of endotoxemia proposed that GDF15 mediated hepatic export of triglycerides was necessary to support the heart during disease (Luan et al., 2019). In our study, we did not find blocking hepatic triglyceride export with the Mttp inhibitor lomitapide to significantly influence survival, and we did not find differences in fatty acid oxidation in hearts of dying or surviving mice based on indirect measurements. However, there are important differences between our study and the study reported by Luan et al that may help explain such discrepancies. First, in our study fluoxetine-treated mice exhibit elevated serum troponin during sepsis while mice with GDF15 are spared from this. This suggests that fluoxetine-mediated effects on the heart are likely different from those in GDF15 mice. Second, the authors investigated GDF15-regulated triglyceride metabolism using an LPS model of sepsis, while we used a co-infection model with live bacteria. Third, the Luan et al study used male mice, while we used female mice for our studies. Finally, potential microbiota differences may possibly contribute to differences. Future work should be undertaken to further explore triglyceride metabolism dynamics during sepsis, and how this relates to cardiac damage and function, and whether these findings are clinically applicable to human patients.

Our study has revealed how a widely available and safe drug, fluoxetine, promotes immunometabolic host-pathogen cooperation to protect from sepsis induced morbidity and mortality. A recent estimate determined the average cost of the research and development required to bring a new therapy to market as over $1.5 billion (Wouters et al., 2020). One of the most cost-effective and quickest strategies to develop new treatment strategies is repurposing already approved drugs for new conditions. Our study provides sufficient rationale to further explore the therapeutic uses of SSRIs during infection, and reveal new and exciting immunometabolic targets during sepsis.

## Acknowledgements

We thank members of the Ayres lab for helpful comments and suggestions. We thank Karina Sanchez and Alexandria Palaferri Schieber for assistance with an infection experiment. This work was supported by NIH awards DPI AI144249 and R01 AI14929, the NOMIS Foundation (J.S.A.), F31 AI169988 (R.M.G.), and T32 GM007240-43 and T32 GM133351 (P.I. Randolph Y. Hampton, Trainee R.M.G.). This work was supported by the Flow Cytometry Core of the Salk Institute with funding from NIH-NCI CCSG: P30CA014195. Open source icons were obtained from flaticon.com.

## Materials and Methods

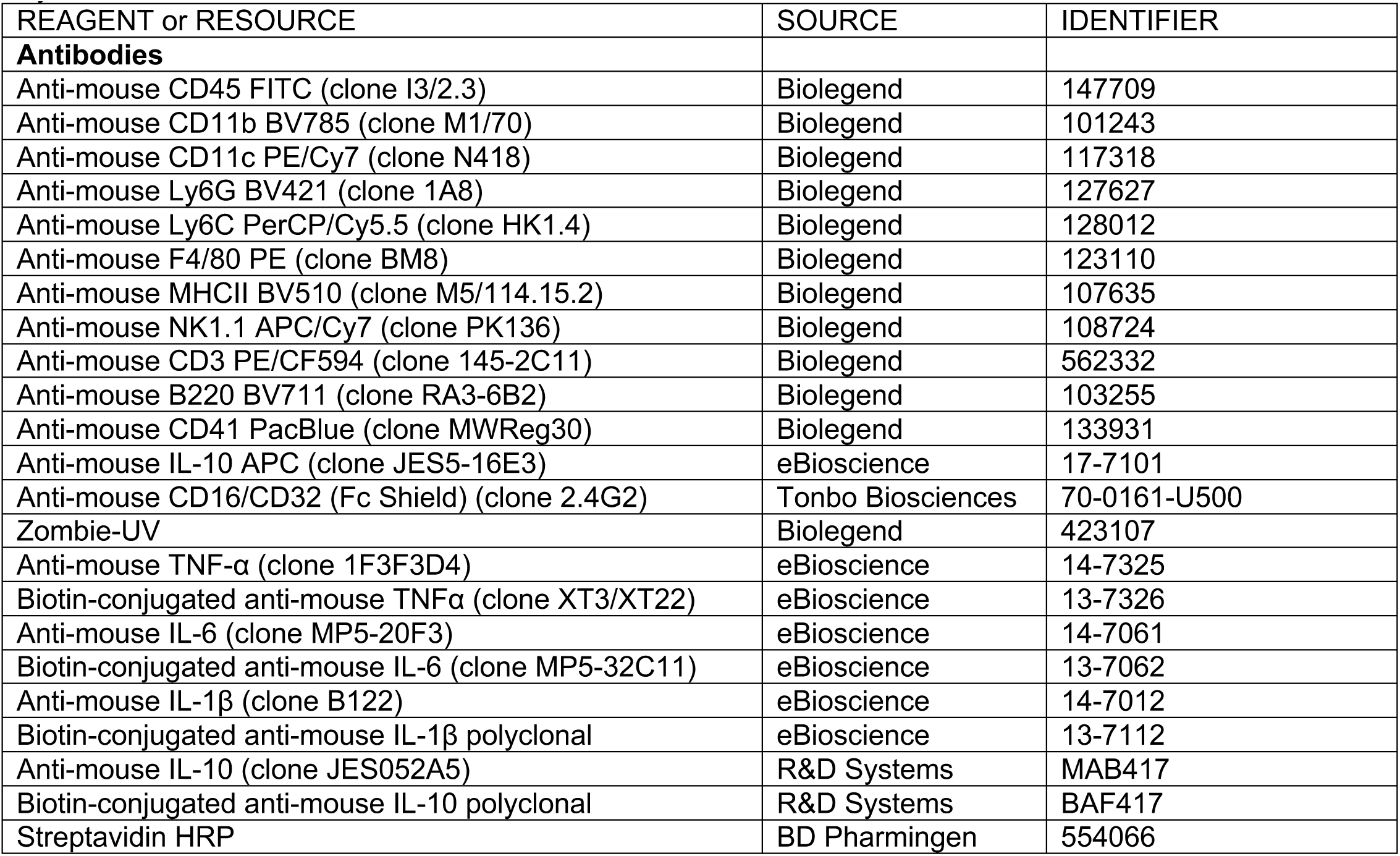
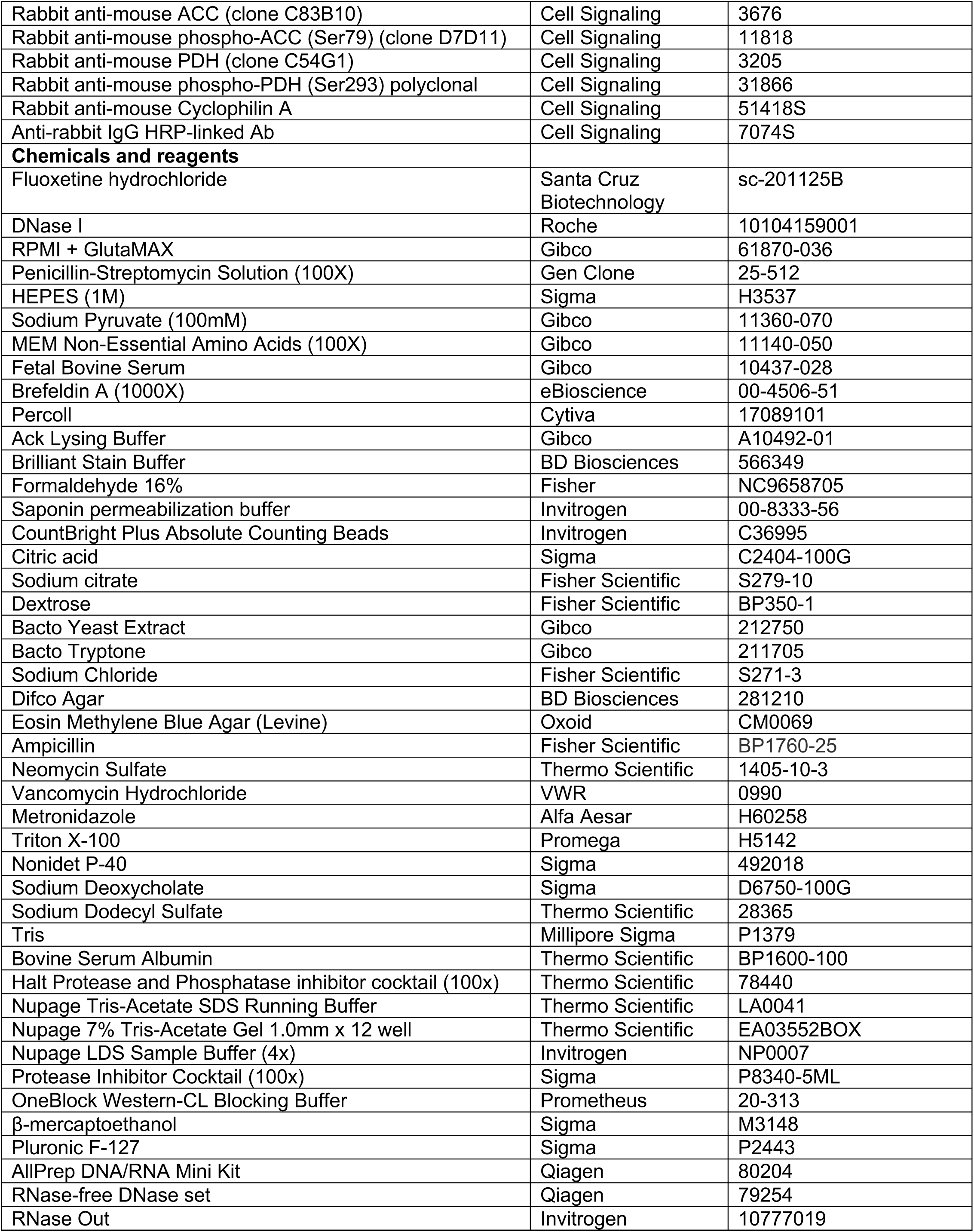
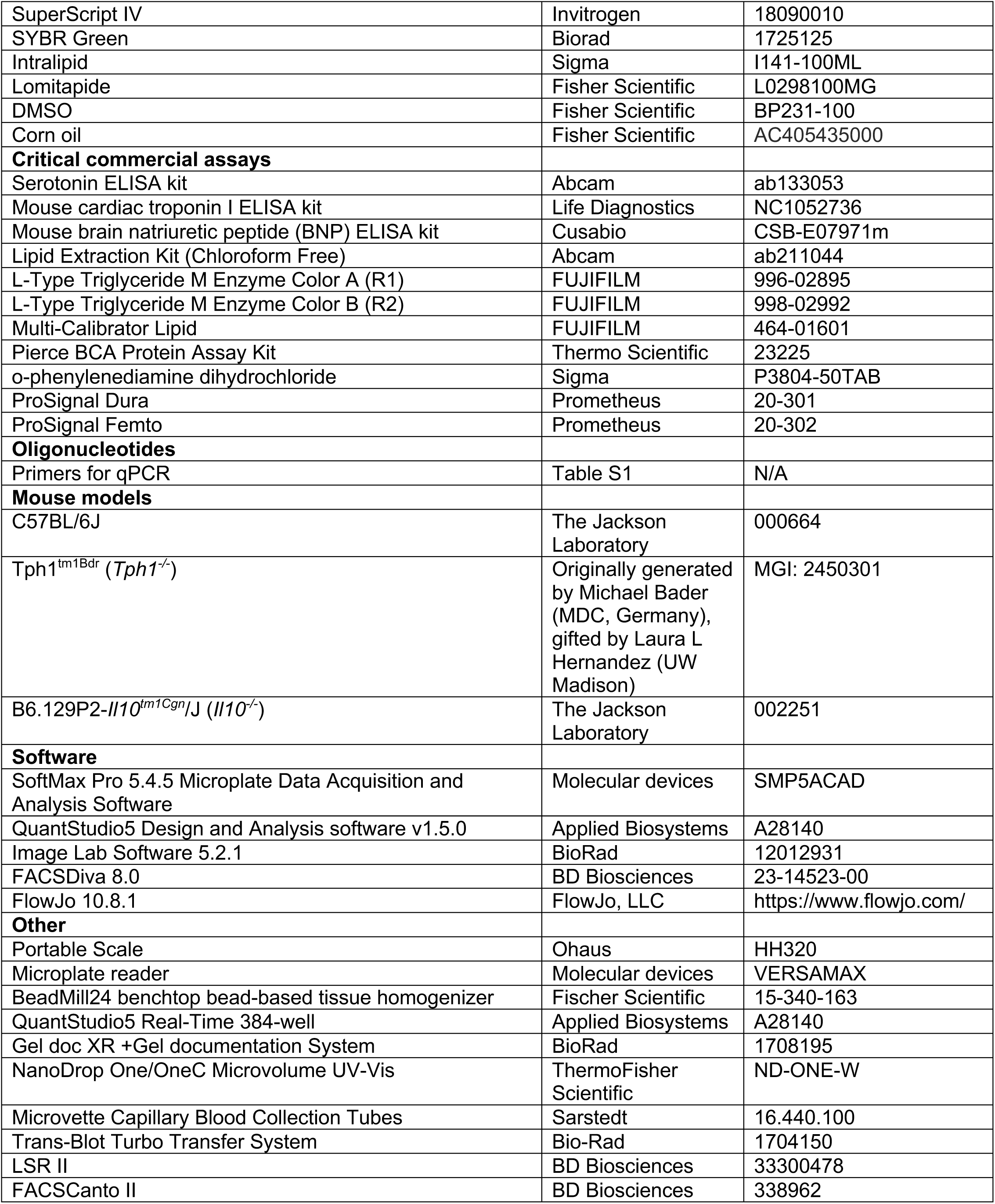
Key Resources Table.

## Mice

Female mice of 10-12 weeks of age purchased from Jackson Labs or bred in our AALAC-certified vivarium were used for the studies described. For experiments with C57BL/6 mice, animals were purchased from Jackson Labs and acclimated in our facility for at least a week prior to experimentation. *Tph1^-/-^*mice (Tph1^tm1Bdr^, MGI:2450301) (Walther et al., 2003) were a generous gift from Dr. Laura L. Hernandez (University of Wisconsin-Madison) and bred in house. *Tph1^-/-^*mice were crossed to C57BL/6 mice ordered from Jackson Labs to generate *Tph1^+/-^*mice. *Tph1^+/-^* were then crossed to each other, and *Tph1^-/-^*and *Tph1^+/+^* littermate females were used for experiments. B6.129P2-*Il10^tm1Cgn^*/J mice were purchased from Jackson Labs and bred in house (Strain #: 002251). Mice were specific pathogen-free, maintained under a 12-hour light/dark cycle, and given standard chow diet *ad libitum* prior to infection. All animal experiments were done in accordance with The Salk Institute Animal Care and Use Committee. In accordance with our IACUC guidelines, in an effort to reduce the number of animals used for our experiments, where possible and appropriate, experiments were done in parallel and shared controls were used. This is indicated in the Figure legends where relevant.

### Bacteria

*Escherichia coli* O21:H+ (Ayres et al., 2012)

*Staphylcoccus aureus* (ATCC strain 12600)

### Culturing *E. coli* O21:H+ and *S. aureus* for mouse infection

*E. coli* O21:H+ was incubated on an EMB plate containing <u>A</u>mpicillin sodium salt (1mg/mL), <u>V</u>ancomycin hydrochloride (0.5mg/mL), <u>N</u>eomycin sulfate (1mg/mL), and <u>M</u>etronidazole (1mg/mL) antibiotics overnight at 37°C to grow single colonies. *S. aureus* was incubated on an LB plate without antibiotics overnight at 37°C to grow single colonies. The next day, a single colony of *E. coli* O21:H+ was inoculated into 100mL LB-AVNM (using antibiotic concentrations noted above) media while a single colony of *S. aureus* was inoculated into 5mL LB without antibiotics. Both cultures were shaken overnight at 37°C (250 RPM). The following morning, the OD600 was measured and an inoculum with a 1:1 mixture of the bacteria was prepped with the indicated doses in sterile 1xPBS that was used directly for mouse infections.

### Fluoxetine treatment

Fluoxetine hydrochloride powder (Santa Cruz Biotechnology) was suspended in 1xPBS at a concentration of 2mg/mL. Mice were administered daily intraperitoneal injections of 10, 20, or 40mg/kg fluoxetine for 7 days pre-infection. 1xPBS was used as vehicle control.

### Mouse infection models

Mice were infected intraperitoneally with 2×10^8^ total bacteria in 500uL delivered through a 25G needle between ZT0 and ZT2 (6:00am – 8:00am). Inoculums were serially diluted and plated to confirm the infectious doses.

Immediately after infection, mice were transferred to a fresh cage and food was removed for the first 10-12 hours post-infection to control for any potential variations in the sickness-induced anorexic response. Mice were clinically monitored as described below every two hours post-infection. Mice that reached clinical endpoints were euthanized according to our animal protocol.

### Survival

Mice were clinically monitored as described below every two hours post-infection. For some experiments, mice were clinically monitored every two hours for the first 10-12 hours post-infection and then again at 24 hours. Mice that had to be euthanized because they reached clinical endpoints during the infection, in addition to those that succumb to the infection, were included in our survival analyses.

### Rectal temperature

Rectal temperatures were taken every two hours post infection for the first 10-12 hours, and then every 24 as noted using the Digisense Type J/K/T thermocouple meter.

### Grading system for monitoring morbidity

We use the following morbidity scale to quantify the morbidity of mice. Infected mice are clinically assessed using this morbidity scale every two hours post-infection. For some experiments, mice were clinically monitored every two hours for the first 10-12 hrs post-infection and then again at 24hrs.

5. Normal. Normal exploratory behavior, rearing on hind limbs, and grooming.
4. Mild. Reduced exploratory behavior, rearing on hind limbs, and grooming. Slower and/or less steady gait, but free ambulation throughout the cage.
3. Moderate. Limited voluntary movement. Slow, unsteady gait for >5 seconds.
2. Severe. No voluntary movement, but mouse can generate slow, unsteady gait for <5 seconds.
1. Moribund. Mouse does not move away from stimulation by researcher and cannot right itself.

### Histology

Livers were harvested and fixed in 10% neutral buffered formalin. Then samples were routinely processed, paraffin embedded, sectioned at 4-5 microns, and hematoxylin and eosin stained. Tissues were evaluated by a board certified veterinary pathologist who was blinded to experimental manipulation, and scored semi-quantitatively for the following parameters: liver necrosis, liver congestion, and hemorrhage. These parameters were scored on a scale of 0-4 with 0 representing normal tissue; 1 represented minimal changes; 2 representing mild changes; 3 representing moderate changes and 4 representing severe changes relative to a score of 1. Representative images were obtained from glass slides using NIS-Elements BR 3.2 64-bit and plated in Adobe Photoshop. Image white balance, lighting, and/or contrast was adjusted using corrections applied to the entire image.

### Cardiac troponin I and BNP quantification

Serum from mice was harvested by cardiac puncture. Troponin I levels were measured using an ultra-sensitive mouse cardiac troponin-I ELISA kit (Life Diagnostics) according to manufacturer’s protocol. BNP levels were measured using (BNP KIT). Plates were read on a VERSAmax microplate reader manufactured by Molecular Devices and data analysis was done using SoftMax Pro.

### BUN, lipase, creatine kinase, ALP, AST, ALT, and quantification

Serums harvested by cardiac puncture were analyzed by IDEXX Bioanalytics.

### Serum triglyceride quantification

As described in the Wako Fujifilm L-Type Triglyceride M kit protocol, 4 μL of serum from tail bleed (for lipid tolerance tests or pluronic tests) or cardiac puncture (for circulating levels during infection) were added to a 96-well plate. Standards were generated by using Wako multicalibrator lipids in the following series (mg/dL): 0, 6, 12, 24, 48, 96, 192, and 384. To each well, 90 μL of R1 was added and the plate was incubated at 37°C for 5 min. The plate was read at 600 nm and 700 nm. After reading, 30 μL of R2 was added to each well and the plate was incubated at 37°C for 5 min. The plate was read again at 600 nm and 700 nm and analyzed following the manufacturer’s instructions. Readings were taken using a 96-well VERSAmax microplate reader and SoftMax Pro software. In Fig. 5A triglyceride measurements were done by IDEXX Laboratories. The protocol utilized by IDEXX Laboratories does not involve an initial glycerol extraction step prior to quantification of triglycerides and therefore the values reported represent both glycerol and triglycerides as noted in the figure. Our in-house triglyceride measurements involve an initial glycerol extraction step prior to quantification of triglycerides and therefore the values reported represent triglyceride levels.

### Quantification of *E. coli* O21:H+ and *S. aureus* in mouse tissues

For quantification of pathogen in organs, colony forming units (CFUs) were quantified. Liver, spleen, kidney, heart, and lung were harvested and homogenized in sterile 1xPBS with 1% Triton X-100 using a BeadMill 24 bench-top bead-based homogenizer (Fisher Scientific). Spleen, kidney, heart, and lung were homogenized in 1 mL while liver was homogenized in 500 µL. Homogenates were serially diluted and plated on LB agar and EMB-AVNM agar and incubated at 37 °C. Colonies were quantified the following day. The limit of detection for all organs was 100 CFUs.

### Bacterial growth curves

*E. coli* and *S. aureus* cultures were grown overnight in the same conditions described for infection inoculum preparation. These cultures were then diluted 1:100 in 200 µL LB in a 96 well plate with varying concentrations of fluoxetine. The plate was grown for 12 hours at 37 °C with shaking and optical density readings were taken at 600 nm every 15 minutes.

### Serotonin quantification

Serum from mice was harvested by cardiac puncture along with liver, spleen, lung, and heart. Plasma was collected by tail bleed into heparin coated microvettes by capillary action. Organs were homogenized in sterile 1xPBS with 1% Triton X-100 using a BeadMill 24 bench-top bead-based homogenizer (Fisher Scientific). Serotonin levels in serum and organ homogenate were measured using a serotonin ELISA kit (Abcam). Plates were read on a VERSAmax microplate reader manufactured by Molecular Devices and data analysis was done using SofMax Pro. Organ serotonin levels were normalized to total organ weight.

### Platelet flow

2 µL of blood was collected by tail bleed throughout infection into microvettes pre-loaded with 2 µL of ACD-A anticoagulant (0.73% citric acid, 2.2% sodium citrate, and 2.45% dextrose). Samples were then immediately transferred into 200 µL of 1xPBS + 2% FBS on ice. Once all samples were collected, cells were washed once with 1xPBS then incubated in 25 µL Brilliant Stain Buffer (BD Biosciences) + extracellular stains for 20 minutes on ice. Cells were then washed in 1xPBS and fixed in 1xPBS + 1%PFA for 10 minutes on ice. Cells were then washed and resuspended in 1xPBS with 5 µL counting beads (Invitrogen) for raw count normalization. Cells were analyzed on a Canto II (BD) using FACSDiva software, and data were analyzed using FlowJo software (Tree Star).

### Cytokine ELISAs

Serum from mice was harvested by cardiac puncture. ELISAs were performed to quantify the levels of IL-1β, IL-6, TNF-α, and IL-10. ELISA antibodies used are listed in the Key Resources Table. Briefly, NUNC 96-well plates were incubated with 50 µL/well primary antibody (final conc. IL-1β 3 µg/mL, IL-6 3 µg/mL, TNF-α 2 µg/mL, IL-10 5µg/mL) in 0.1M sodium phosphate pH 8.0 at 4°C overnight, washed three times with 1xPBS+0.1%Tween20, and then blocked for 1 hour at 37°C with 100 µL/well 1xPBS+1%TritonX-100+1%BSA. After washing once, 50 µL of samples and standards (diluted in 1xPBS+1%TritonX-100) were loaded and incubated at 37°C for 1 hour. Following four washes, 50 µL/well secondary antibody (final conc. IL-1β 500 ng/mL, IL-6 500 ng/mL, TNF-α 1 µg/mL, IL-10 500 ng/mL) was added in 1xPBS+1%TritonX-100+1%BSA and the plate was incubated for 30 minutes at 37°C. After washing three times, 50 µL/well of 1:1000 Streptavidin HRP in 1xPBS+1%TritonX-100+1%BSA was added, and the plate was incubated for 15 minutes at 37°C. Following three more washes, 50 µL/well of an OPD-based developing solution (5mL citrate buffer [0.1M sodium phosphate, 0.02M citric acid], 1 tablet of OPD, and 2 µL of 30% H_2_O_2_) was applied to the plate and the reaction was stopped after 15 min at room temperature using 20 µL 3M HCl. The optical density at 490 nm was determined using a SpectraMax (Molecular Devices) plate reader and analyzed using SoftMax Pro software 5.4.

### qRT-PCR

Livers and hearts were harvested and snap frozen in liquid nitrogen then stored at –80°C. Frozen tissues were ground into a powder using a mortar and pestle chilled in liquid nitrogen. RNA was isolated from ∼20mg tissue powder using Qiagen Allprep DNA/RNA Mini Kit. Briefly, the sample was lysed in 600 µL RLT buffer containing 10 µL β-mercaptoethanol using a BeadMill 24 bench-top bead-based homogenizer (Fisher Scientific) and centrifuged for 3 minutes at 18,000g in a microcentrifuge. The supernatant was transferred to an AllPrep DNA column and centrifuged for 30 s at 18,000g. Flow-through was mixed with 70% ethanol and transferred to an RNeasy spin column and centrifuged for 30 seconds. The flow-through was discarded and the column containing RNA was washed with 350 µL RW1 buffer. Flow-through was discarded. The column was treated with Qiagen RNase-free DNase1 (10 µL DNase I [2.73 Kunitz units / µL] and 70 µL buffer RDD) for 15 minutes at room temperature. The column was washed again with 350 µL RW1 and the flow-through was discarded.

The column was then washed with 500 µL of RPE buffer. The flow-through was discarded and the column was spun again to remove any residual wash buffer. After the final spin, the column was placed in a new Eppendorf tube, and RNA was eluted with 40 µl of RNase-free water. cDNA was made using the SuperScript IV kit (Invitrogen) after all samples were diluted to 20 ng/µL. Real-time qPCR was performed using iTaq SYBR Green Mix (BioRad) on a ThermoFisher QuantStudio 5 qPCR machine. Relative standard curves were generated by mixing cDNA from each sample and then making serial dilutions of the mix. Analysis of gene expression was then done by comparing Ct of the gene of interest to the relative standard curve for that gene, then normalizing this to the expression of the housekeeping gene (*rps17*) for each sample. See table 1 for primer sequences.

### Peritoneal lavage flow cytometry preparation

Peritoneal lavage was collected immediately after euthanasia. Briefly, the mouse skin was peeled back to expose the peritoneal cavity. 5mL of 1xPBS was injected into the peritoneal cavity, taking care not to puncture any internal organs. The mouse was then shaken for 10 seconds, then the lavage fluid was taken back up into the syringe. The lavage fluid was then mixed with 10mL of RPMI complete media (RPMI GlutaMAX, 1% Pen/Strep, 10% heat inactivated FBS, 1% HEPES 1M, 1% sodium pyruvate, 1% MEM-NEAA) on ice. Once all samples were collected, they were centrifuged at 400g for 4 minutes and the supernatant was poured off. 2 mL of Ack lysis buffer was added and samples were incubated for 2 minutes before 10 mL of RPMI complete media was added. Samples were again centrifuged at 400g for 4 minutes. Supernatant was discarded and samples were transferred to a 96-well v-bottom plate.

### Flow cytometry cell incubation, staining, and analysis

Samples in a 96-well v-bottom plate were resuspended in 200 µL RPMI complete media + 1:1000 Brefeldin A. The plate was incubated at 37 °C and 5% CO_2_ for 4 hours. Following incubation, cells were washed once in 1xPBS then treated with 25 µL 1xPBS + 1:100 Fc block + 1:100 Zombie UV for 10 minutes on ice. Cells were then washed in 1xPBS then incubated in 25 µL Brilliant Stain Buffer (BD Biosciences) + extracellular stains for 20 minutes on ice. Cells were then washed in 1xPBS then fixed in 200 µL 1% formaldehyde overnight at 4°C. The following day cells were washed with 1xPBS and stained for 30 minutes with intracellular stains in 50 µL saponin permeabilization buffer (Invitrogen). The cells were then washed again and resuspended in 200 µL PBS. 5 µL counting beads (Invitrogen) were added to each sample for raw count normalization. Cells were analyzed on a LSR II (BD) using FACSDiva software, and data were analyzed using FlowJo software (Tree Star).

### Heart triglyceride measurement

Hearts were removed from the body cavity, blood was drained from the heart chambers, and then the organs were snap frozen in liquid nitrogen. Frozen organs were ground into a powder using a mortar and pestle chilled in liquid nitrogen. Lipids were extracted from ∼20mg organ powder using a lipid extraction kit (Abcam). Briefly, powder was homogenized in 375 µL of extraction buffer using a BeadMill 24 bench-top bead-based homogenizer (Fisher Scientific). The homogenate was then transferred to a fresh 1.5mL Eppendorf tube and centrifuged at 10,000g for 5 minutes at 4°C. 250 µL was then transferred into a PCR strip. The samples were then allowed to evaporate overnight at 37°C. The following day samples were resuspended in 50 µL suspension buffer by vigorous sonication and vortexing. Triglycerides were measured using Fujifilm L-Type Triglyceride M kit, following the manufacturer’s protocol.

### Western blot

Heart tissue powder was homogenized in 600 µL RIPA buffer supplemented with 1:100 Protease Inhibitor Cocktail (Sigma) and 1:100 Halt Protease and Phosphatase Inhibitor Cocktail (Thermo Scientific). Lysates were centrifuged at 4°C for 30 minutes at 18,000g and transferred to a new tube. Protein concentration was quantified with a BCA reaction. Samples were diluted in RIPA + a 1:10 mixture of 2-mercaptoethanol and NuPAGE LDS Sample Buffer (4X) (Invitrogen) to a final protein concentration of 1µg/µL, then incubated at 70°C for 10 minutes and sonicated in a water bath for 10 minutes. 15 µL of each sample was loaded into 7% NuPage 1.0mm x 12 well Tris-Acetate gels with Tris-Acetate SDS running buffer (50mL 20X Tris Acetate SDS in 950 mL DI H_2_O for 60 minutes at 150V. Gels were placed on Trans-Blot® Turbo™ Midi 0.2 µm Nitrocellulose Transfer Packs (Bio-Rad) and transferred in Trans-Blot® Turbo™ Transfer System (Bio-Rad) (2.5A 25V for 10 minutes). Membranes were stained with Ponceau Total Protein Stain (Prometheus) to visualize total protein, then membranes were cut into strips for each protein to be assayed. The strips were then washed with 1X TBST to de-stain, then placed into blocking solution (5% BSA in 1X TBST) on a shaker platform for one hour at room temperature. Blocking solution was removed and primary antibody solutions were added to membranes (1:1000 Ab in Prometheus OneBlock™ Western-CL Blocking Buffer) and placed on shaker platform overnight at 4°C. Primary antibody solutions were removed and membranes were washed on shaker platform for 5 min with 1X TBST, repeated 4 times at RT. Secondary antibody solution was added containing 1:3000 anti-rabbit IgG HRP-linked Ab in blocking buffer (5% BSA in 1X TBST) and membranes were placed on shaker platform for 1hr at RT. Secondary antibody solutions were removed and membranes were washed again on a shaker platform for 5 min with 1X TBST, repeated 4 times at RT. Nitrocellulose blots were developed using a mixture of Femto and Dura chemiluminescent reaction and visualized with a Bio-Rad Gel Doc XR+ machine. For blots targeting total and phosphorylated forms of the same protein, the same lysate was run on two different gels at the same time in the same gel rig. The gels were then transferred to the same membrane. After staining of the membrane with Ponceau, the membranes were cut and destained. The cut membranes were then probed for the relevant protein. Total and phosphorylated blots had their own Cyclophilin A loading control that was run on the same gel.

### Lipid tolerance test

Mice were retro-orbitally injected with 100 µL Intralipid (Sigma 20% emulsion) at a dose of 10 mL/g body weight 7 hours post-infection. Blood was collected via the lateral tail vein at 0, 15-, 30-, 45-, and 60-minutes post-injection. Blood was centrifuged at 6,000 rpm for 20 min and serum was collected for triglyceride measurements. Total triglycerides were measured using Fujifilm L-Type Triglyceride M kit, following the manufacturer’s protocol.

### Pluronic test

To make pluronic solution, 5 g of pluronic (Sigma Pluronic F-127) was dissolved in 50 mL of water and the solution was filter sterilized with a 0.22 µm filter. Mice were intraperitoneally injected with the pluronic solution at a dose of 10 mL/g of bodyweight at the time of infection. Blood was collected by tail vein at 0, 60-, 120-, 180-, and 240-minutes post-injection. Blood was spun at 6,000 rpm for 20 min and serum was collected for triglyceride measurements. Triglycerides were measured using Fujifilm L-Type Triglyceride M kit, following the manufacturer’s protocol.

### Lomitapide treatment

Mice were orally gavaged with 10mg/kg lomitapide in 2% DMSO + corn oil, or administered 2% DMSO + corn oil vehicle, daily for three days pre-infection.

### Statistical analysis

Statistical tests were done using Prism version 8.4.3. Sample sizes, number of independent experiments pooled, statistical tests used, and p-values for each figure and panel are indicated in the figure legends.

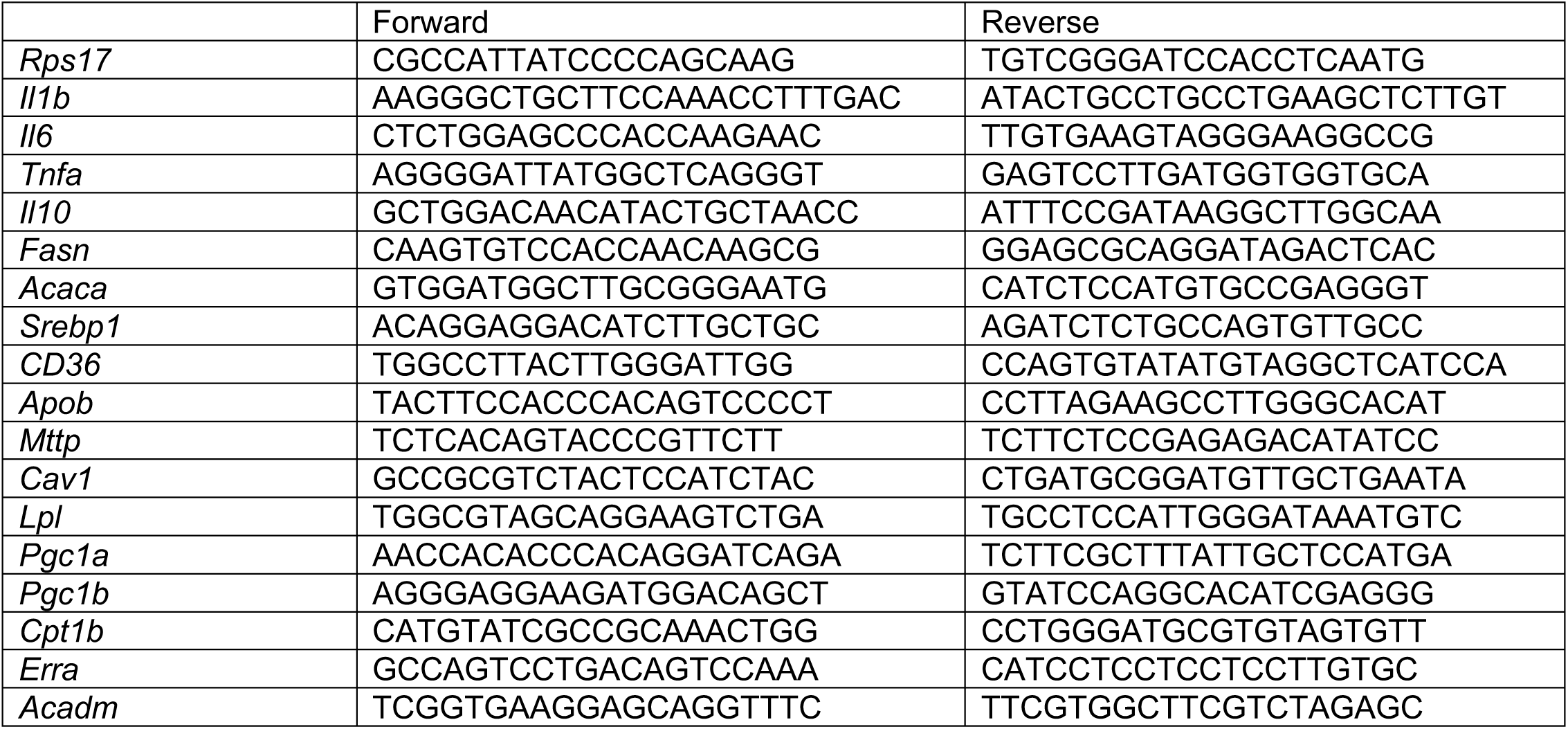
Table S1.

## Notes

### Competing Interest Statement

The authors have declared no competing interest.

