## Supplemental materials for "Fluoxetine promotes immunometabolic defenses to mediate host-pathogen cooperation during sepsis"

**Supplemental Figure Legends**

**Supplemental Figure 1: Fluoxetine protects from polymicrobial sepsis.**
(A-C) Histological analysis of H&E stained livers at 10 hours post infection in vehicle or fluoxetine treated mice infected with polymicrobial sepsis. n = 5 per condition, one independent experiment shown. Data represent mean ± SEM. Two-way ANOVAs with Tukey's multiple comparisons.
(D-E) *E. coli* or *S. aureus* burden analysis from vehicle or fluoxetine treated mice at 8-10 hours post infection. n = 15 per condition, three independent experiments combined. Data represent geometric mean ± geometric SD. Unpaired t-tests.
(F-G) *E. coli* or *S. aureus* growth curves grown in varying concentrations of fluoxetine. n = 4 per condition, one independent experiment.
(H-Q) Reaction norm analyses plotting body temperature at the time of dissection against *E. coli* or *S. aureus* CFUs. (H) *E. coli* liver, (I) *E. coli* spleen, (J) *E. coli* kidney, (K) *E. coli* heart, (L) *E. coli* lung, (M) *S. aureus* liver, (N) *S. aureus* spleen, (O) *S. aureus* kidney, (P) *S. aureus* heart, (Q) *S. aureus* lung. Same mice as in (A-B). Semilog linear regression, y-intercept constrained to uninfected temperature, Extra sum-of-squares F Test to compare slopes.
* $p<0.05$, ** $p<0.01$, *** $p<0.001$.

**Supplemental Figure 2: Fluoxetine regulates the degree and duration of the inflammatory response during sepsis.**
(A-B) (A) Plasma serotonin levels throughout infection of mice treated with vehicle or fluoxetine infected with polymicrobial sepsis. (B) Area under the curve analysis of (A). n = 5 per condition, one independent experiment. For time-course, two-way ANOVA with Dunnett's multiple comparisons. For AUC, unpaired t-test.
(C-D) (C) Gating strategy and (D) Platelet count at 8 hours post infection of mice treated with vehicle or fluoxetine infected with polymicrobial sepsis. n = 5 per condition, one independent experiment. Two-way ANOVA with Tukey's multiple comparisons.
(E-H) Hepatic transcript levels of (E) *Tnfa*, (F) *Il6*, (G) *Il1b*, and (H) *Il10* in vehicle or fluoxetine treated mice infected with polymicrobial sepsis. n=5-15 per condition, three independent experiments combined. Unpaired t-tests with Holm-Sidak multiple comparisons correction.
(I) Percentage of indicated population that was IL10+, same mice as panel 4E. Unpaired t-tests with Holm-Sidak multiple comparisons correction.
(J-K) Gating strategies for panel 4E.
(L-N) Hepatic transcript levels of (K) *Tnfa*, (L) *Il6*, and (M) *Il1b* at 10 hours post infection in wildtype or *Il10*$^{-/-}$ mice pre-treated with fluoxetine infected with polymicrobial sepsis. n=5-11 per condition, two independent experiments combined. Two-way ANOVA with Tukey's multiple comparisons test.
In all panels data represent mean ± SEM. * $p<0.05$, ** $p<0.01$, *** $p<0.001$.

**Supplemental figure 3: IL-10 is required for fluoxetine-mediated cooperative defenses during sepsis.**
(A-B) (A) *E. coli* and (B) *S. aureus* burden analysis at 10 hours post infection from wildtype or *Il10*$^{-/-}$ mice treated with fluoxetine infected with polymicrobial sepsis. Data represent geometric mean ± geometric SD. Same mice as in panel 4L. Unpaired t-tests.
(C-G) Reaction norm analyses plotting body temperature at the time of dissection against total, *E. coli*, or *S. aureus* CFUs. (C) Liver, (D) Spleen, (E) Kidney, (F) Heart, (G) Lung. Same mice as in panel 4L. Semilog linear regression, y-intercept constrained to uninfected temperature, Extra sum-of-squares F Test to compare slopes.

**Supplemental figure 4: Lpl activity is required for fluoxetine-mediated defenses during sepsis.**
(A) Survival of vehicle and lomitapide treated mice infected with polymicrobial sepsis. n=10 per condition, one independent experiment shown. Log-rank analysis.
(B-C) (B) *E. coli* or (C) *S. aureus* burden analysis at 8-10 hours post infection in fluoxetine treated mice with or without Pluronic F-127 injection at the time of infection with polymicrobial sepsis. Data represent geometric mean ± geometric SD. Same mice as in panel 5I. n=10 per condition, two independent experiments combined. Unpaired t-tests.

(D-H) Reaction norm analyses plotting body temperature at the time of dissection against total, *E. coli*, or *S. aureus* CFUs. (D) Liver, (E) Spleen, (F) Kidney, (G) Heart, (H) Lung. Same mice as in panel 5I. Semilog linear regression, y-intercept constrained to uninfected temperature, Extra sum-of-squares F Test to compare slopes. * $p<0.05$.

**Supplemental Figure 5: Uncropped cardiac western blots.**
(A) Uncropped western blots from Figure 7A.
(B) Uncropped western blots from Figure 7D.
(C) Uncropped western blots from Figure 7H.
(D) Uncropped western blots from Figure 7K.

### Supplemental Figure 1

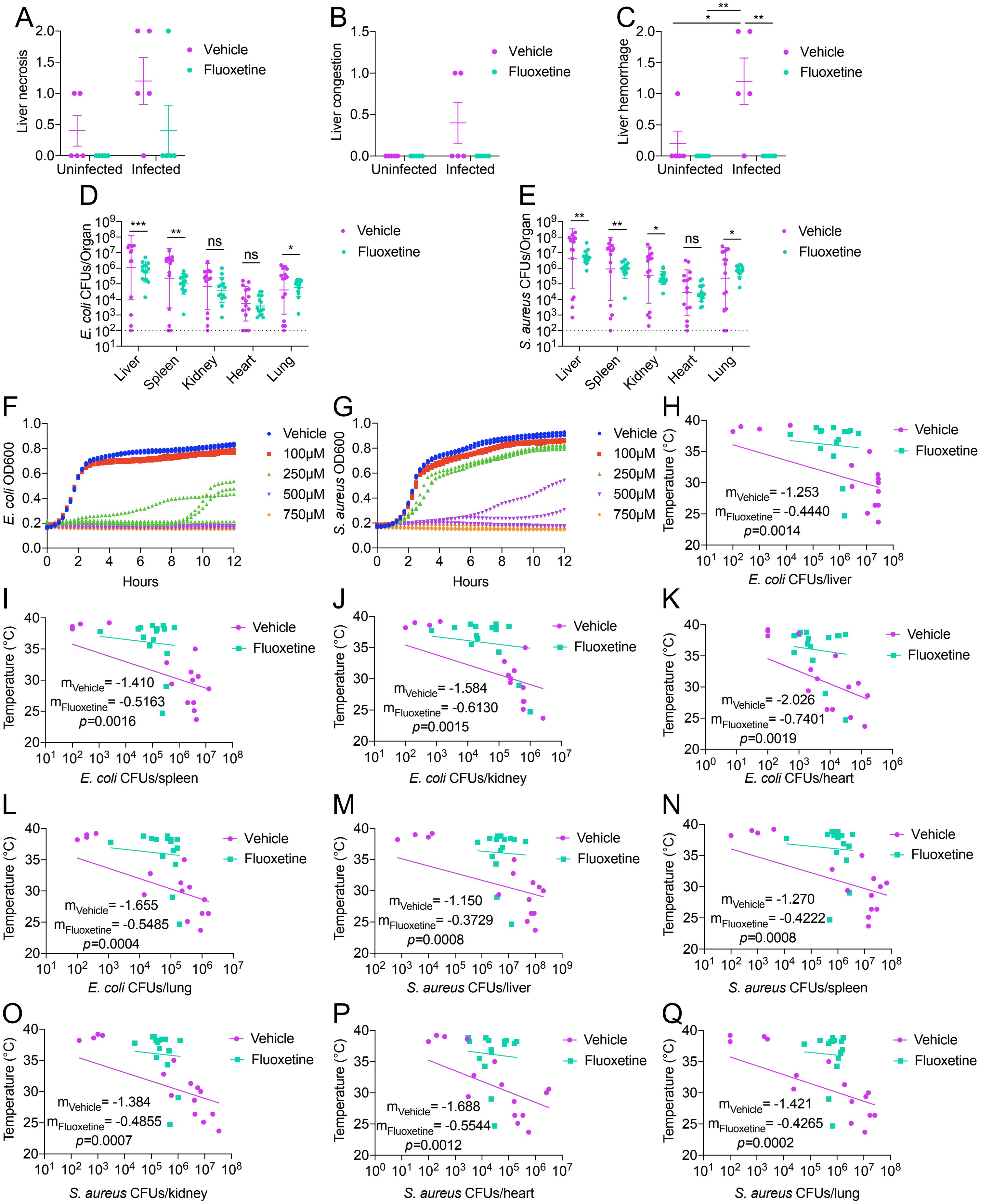

### Supplemental Figure 2

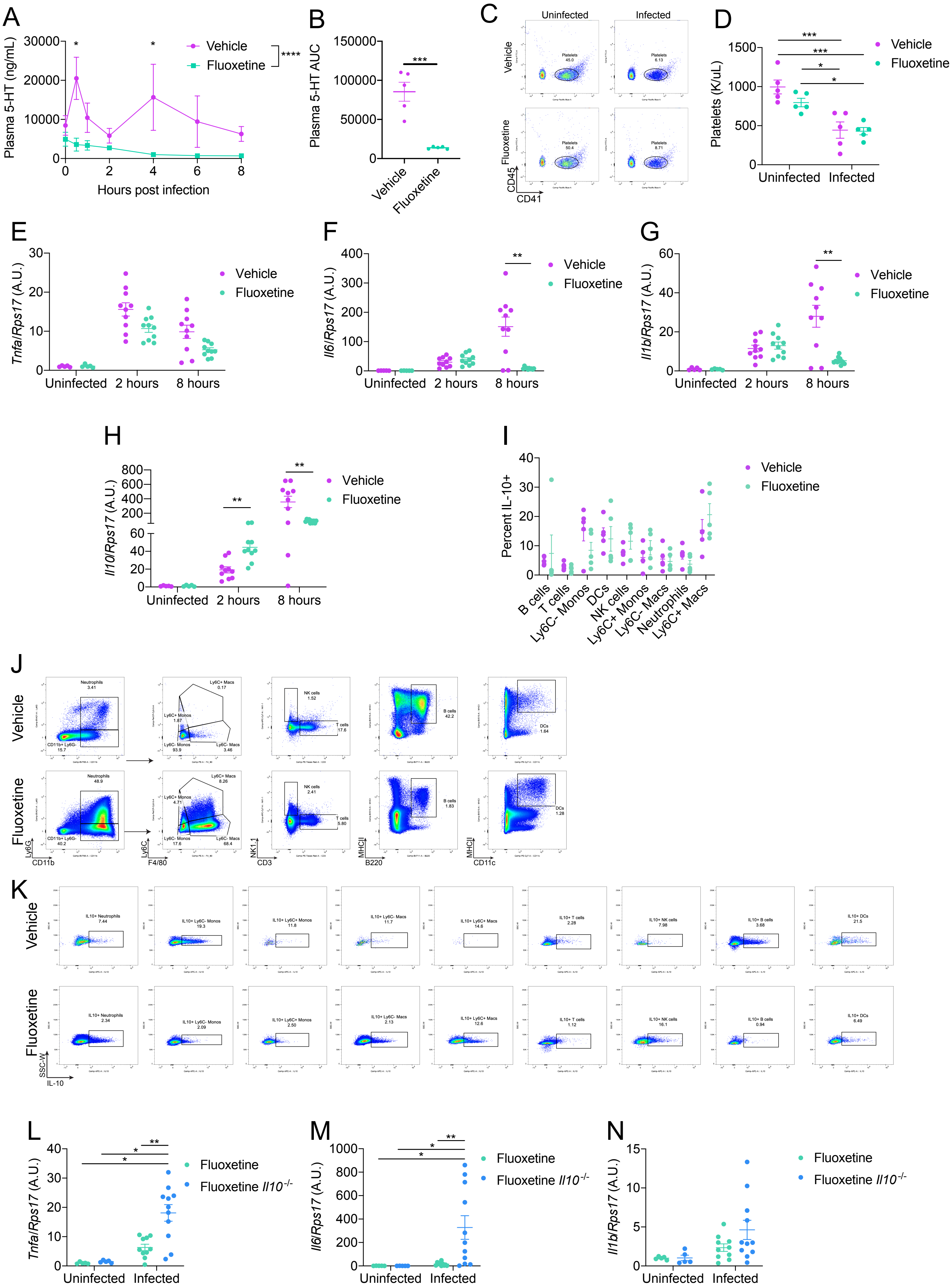

### Supplemental Figure 3

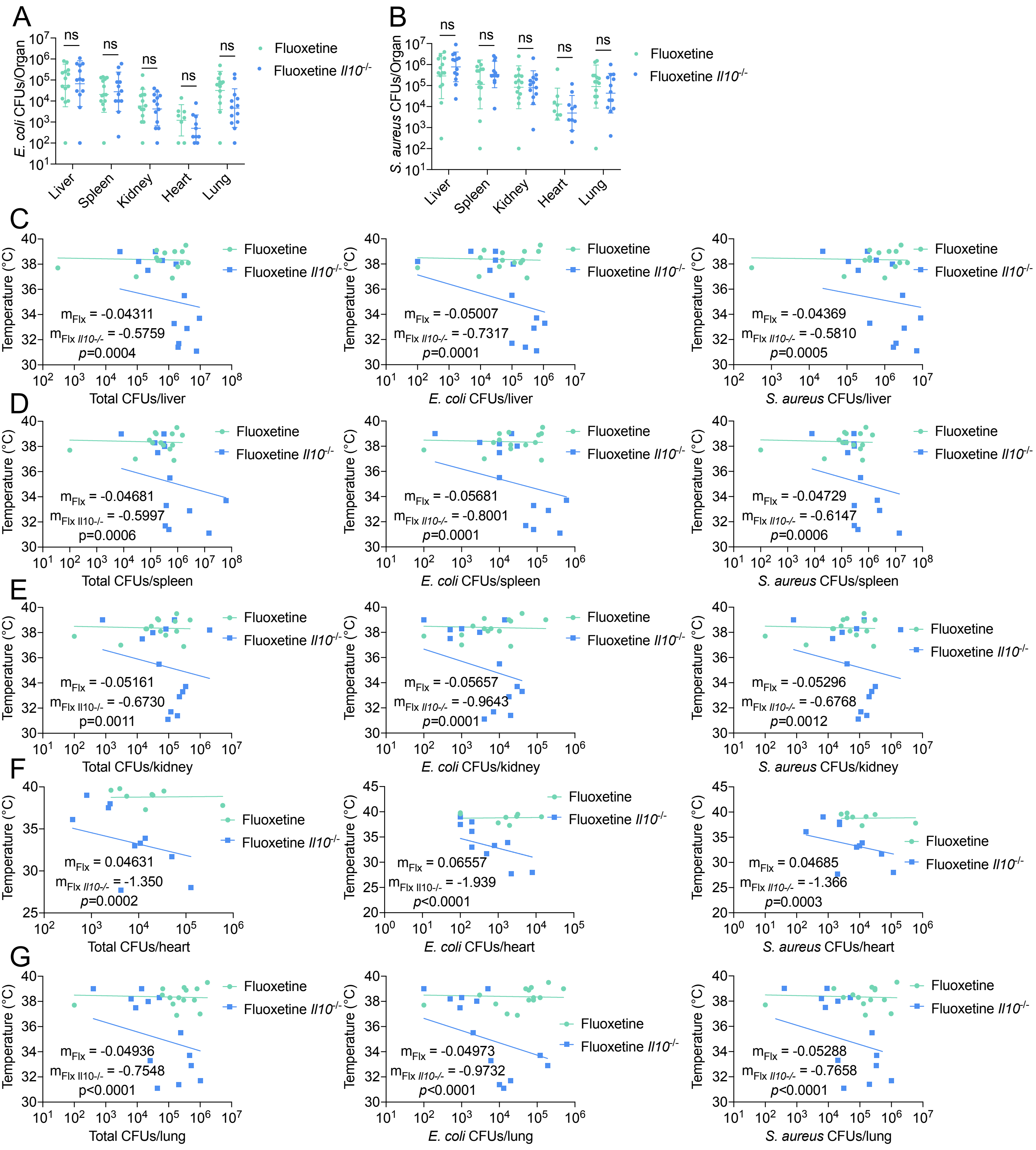

### Supplemental Figure 4

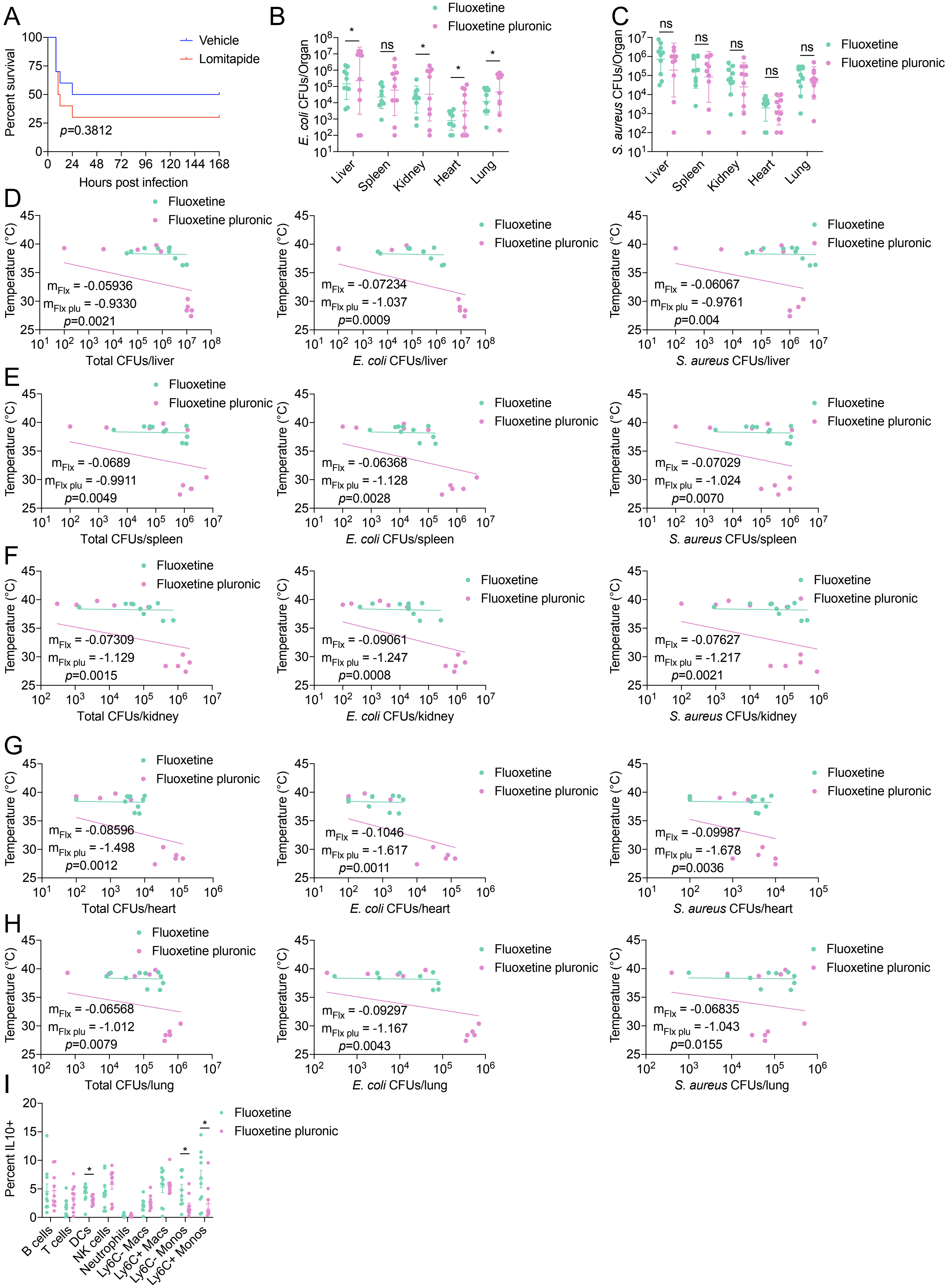

### Supplemental Figure 5

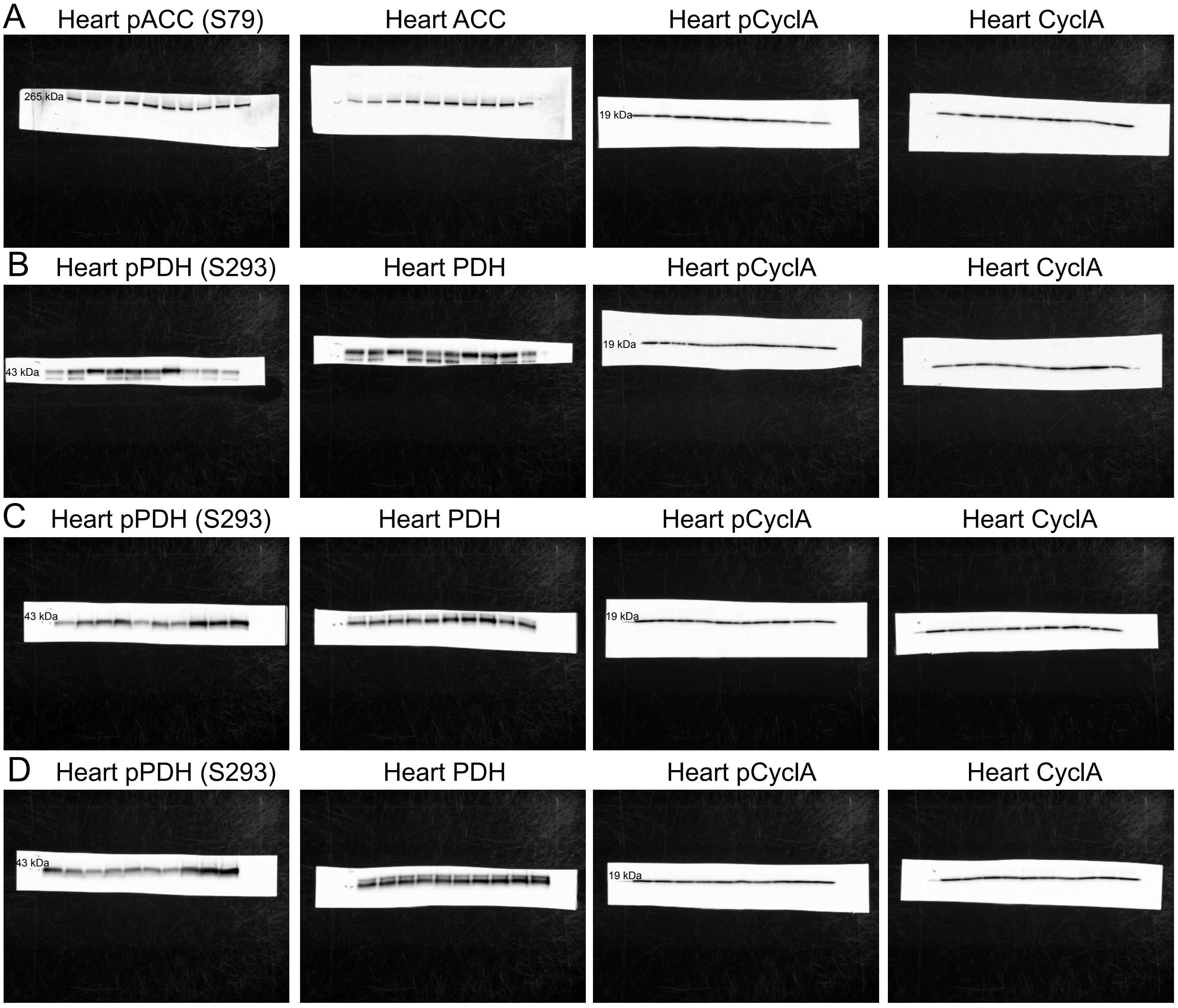
